# Fetal MRI Reveals Altered Subplate Development in Congenital Heart Disease

**DOI:** 10.64898/2026.08.18.745587

**Authors:** Andrea Gondová, Seungyoon Jeong, Nicole Stepovich, Wayne Tworetzky, Victoria R. Bradford, Anjali Sadhwani, Jennings Zhang, Sungmin You, P. Ellen Grant, Kiho Im, Caitlin K Rollins

## Abstract

**Background:** The subplate is a transient fetal brain compartment that provides an early foundation for downstream cerebral development. Congenital heart disease (CHD) alters fetal circulation and cerebral substrate delivery, but its impact on subplate development and whether resulting alterations relate to later neurodevelopmental outcomes remain unclear.

**Methods:** In this retrospective observational cohort study, We used fetal MRI to quantify whole-brain, lobar, and regional (17 bilateral cortical regions) subplate volume and thickness to evaluate group differences between 76 fetuses with CHD and 62 typically developing (TD) fetuses scanned between 21-32 weeks of gestation, and estimated individualized deviations from TD developmental trajectories. Associations with fetal hemodynamic indices (substrate delivery score, cerebroplacental ratio [CPR]) and two-year neurodevelopmental outcomes (Bayley Scales of Infant and Toddler Development, N=46 CHD, N=37 TD) were explored.

**Results:** Whole-brain subplate volume was lower in CHD, corresponding to a 5.7% reduction relative to age- and sex-expected values (p=0.003), but this difference was substantially attenuated after accounting for global brain volume (p=0.090). In contrast, regional analyses identified persistent spatially structured deviations beyond global scaling, most consistently involving posterior parietal, occipital and temporal regions, with a left-hemisphere bias in subplate thickness. Normative modelling demonstrated bidirectional regional deviations and increased inter-individual variability in CHD, with extreme subplate volume deviations enriched across 73% of cortical regions (p=0.009). Higher CPR was associated with lower SP thickness deviations, with the association strengthening after accounting for cerebral substrate delivery (p=0.010), although these analyses were exploratory. In CHD fetuses with postnatal follow-up, prenatal subplate deviations showed modest associations with neurodevelopmental outcomes, with right precuneus subplate thickness associated with receptive (β=-11.87, q=0.017) and expressive (β=-14.50, q=0.039) communication on the Bayley, after correction for multiple comparisons.

**Conclusions:** Fetal subplate alterations in CHD are dominated by global reductions in brain growth but also include spatially heterogeneous and individually variable regional deviations beyond global scaling. Exploratory associations with fetal hemodynamics and postnatal neurodevelopment provide hypotheses for future studies investigating the developmental significance of these prenatal alterations.

## 1. Introduction

Congenital heart disease (CHD) is the most prevalent major birth defect in the United States^1^, affecting approximately 1% of live births. Despite substantial advances in surgical and perioperative care, up to 50% of patients with severe CHD suffer from long-term neurodevelopmental impairments.^2^ Increasing evidence suggests that these alterations originate prenatally.^3^ Fetal MRI studies have demonstrated that fetuses with CHD diverge from typically developing (TD) cohorts across multiple aspects of brain development, including reduced global and regional brain volumes^4–9^, altered cortical surface expansion^10,11^, delayed gyrification^12,13^, microstructural abnormalities^14^, and alterations in structural connectivity and myelination^12,13,15,16^. Additional studies have identified alterations in brain metabolism^7^, tissue oxygenation, and cerebral oxygen consumption^17,18^, and recent reports suggest associations between prenatal brain alterations and later neurodevelopmental outcomes^10,19,20^.

The mechanisms underlying altered fetal brain development in CHD are complex and multifactorial, involving genetic contributions^21^, placental dysfunction^22,23^, and altered fetal systemic and cerebral hemodynamics that may impair cerebral delivery of oxygen and metabolic substrates, particularly glucose.^7,18,24,25^ The fetal brain is among the most metabolically active organs, relying heavily on placental oxygen and glucose supply to support rapid developmental processes.^26^ Consequently, both impaired cerebral substrate delivery and altered cerebrovascular hemodynamics have been proposed as potential contributors to impaired cerebral development in CHD, including through chronic cerebral hypoxia and reduced metabolic substrate availability.^18,24^ In response, the fetal cerebral circulation can mount compensatory adaptations, including reduced cerebrovascular resistance and vasodilation aimed at preserving cerebral blood flow (“brain sparing”), a well-characterized response to hypoxic stress that can be assessed using Doppler ultrasound.^27,28^ For example, recent work suggests that in forms of CHD with the most impaired cerebral perfusion (e.g. systemic outflow tract obstruction and/or single ventricle physiology), brain sparing, assessed by lowered cerebroplacental ratio and middle cerebral artery pulsatility index, is associated with improved neurodevelopmental outcomes.^29^ Nevertheless, substantial heterogeneity exists in the literature regarding this relationship, and some Doppler studies suggest that compensatory mechanisms may be inadequate in more severe or sustained circulatory compromise.^30^ Importantly, these physiological disturbances are unlikely to affect cortical development uniformly, but may instead interact with regionally heterogeneous maturational timing, connectivity, and metabolic demand across the developing brain.

One transient developmental compartment that may be particularly susceptible to such disturbances is the subplate (SP). The SP undergoes rapid expansion during the 2^nd^ and 3^rd^ trimesters and represents one of the most metabolically active compartments of the developing brain, playing a central role in early cortical development, serving as a substrate for thalamocortical and corticocortical connectivity, neuronal migration, axonal guidance, and early circuit formation.^31^ Additionally, typical SP development exhibits marked spatial and temporal heterogeneity^32^, suggesting that developmental disturbances may likewise show regionally specific patterns of vulnerability. Experimental and neuropathological studies suggest that CHD-related brain injury is associated with immature cortical development and white matter abnormalities, implicating vulnerability of subplate-dependent developmental processes.^33,34^ Additionally, SP neurons are particularly sensitive to hypoxic-ischemic injury and disturbances in cerebral perfusion during periods of rapid early circuit formation.^35–37^ Indeed, recent fetal brain MRI studies demonstrate reduced global and lobar SP volume and thickness, especially in fetuses with hypoplastic left heart syndrome (HLHS) and transposition of the great arteries (TGA) compared with healthy controls.^4,8^ Altered SP maturation has been previously hypothesized in several neurodevelopmental disorders^38,39^ and studying its alterations in CHD may therefore represent an early window linking developmental disruption and later cognitive, motor, and behavioral outcomes.

The extent to which SP alterations in CHD reflect generalized reductions in overall brain growth versus disproportionate regional abnormalities remains unclear. Distinguishing these effects may be particularly important for understanding selective cerebral vulnerability in CHD. Moreover, substantial heterogeneity exists across CHD phenotypes and across individuals, raising the possibility that altered SP development may follow spatially heterogeneous and individualized trajectories rather than reflecting a uniform pattern of injury. Here, we investigated SP development in fetuses with CHD scanned prior to 32 weeks’ gestation using fetal MRI-based SP segmentation, surface-based SP thickness analysis, and TD-referenced normative developmental modelling. We characterized SP volume and thickness at whole-brain, lobar, and regional scales and examined whether regional alterations persisted after accounting for overall brain growth. In addition, we quantified individualized deviations from TD developmental trajectories to assess spatial heterogeneity and inter-individual variability in SP development. We subsequently explored whether SP developmental deviations were associated with cerebral substrate delivery and cerebroplacental hemodynamics, representing complementary aspects of the altered fetal circulatory environment in CHD, as a first step toward understanding potential mechanistic contributors. Finally, we evaluated how SP deviations relate to postnatal neurodevelopmental outcomes. We hypothesized that fetuses with CHD would demonstrate spatially heterogeneous alterations in SP development, reflecting regionally selective vulnerabilities exceeding global reductions in brain growth.

## 2. Materials and Methods

### 2.1. Cohort characteristics

This study was conducted under a protocol approved by the Boston Children’s Hospital Institutional Review Board (IRB-P00008836), and written informed consent was obtained from pregnant women/parents of infants. Eligibility criteria were previously reported.^8^ Cohort consisted of CHD fetuses and control group of fetuses with a family history of CHD but normal fetal and postnatal echocardiographic findings recruited from 2014 to 2025. Fetuses with incidental extra-cardiac, neurological, or genetic abnormalities identified after enrollment were excluded, as were subjects with insufficient MRI quality (SI-Fig1). Only fetuses scanned before 32 weeks’ gestational age (wGA) were included because SP contrast becomes indistinguishable from the intermediate zone on T2-weighted MRI beyond this timepoint (SI-Fig2).

Structural and hemodynamic data from fetal echocardiography and Doppler assessment were obtained via standard institutional clinical protocols. Measures included pulsatility index (PI) in the middle cerebral artery (MCA) and umbilical artery (UA). Lower PI values reflect decreased pulsatility with increased diastolic flow, indicating lower vascular resistance. ^29^ From these, we calculated the cerebroplacental ratio (CPR) as the ratio of MCA PI to UA PI. By integrating changes in both measures, CPR provides a composite index of relative cerebroplacental hemodynamics that captures both placental circulatory insufficiency and adaptive changes in the fetal cerebral circulation.^40^ CPR was standardized into Z-scores according to.^41^ CPR values <1 were considered abnormal, consistent with prior studies.^28^ When multiple fetal assessments were available, the examination acquired most closely in time to the MRI session was selected, and the delay between echocardiography and MRI acquisition (ECHO-MRI delay) was included as a covariate in analyses including CPR.

For the CHD group, substrate delivery scores were calculated by summing three key physiological and anatomic features on fetal ECHO as previously described^8^: aortic arch flow direction (antegrade=0; retrograde=1), ventricular physiology (two-ventricle [2V]=0; single-ventricle [SV]=1), and substrate concentration (normal=0; mixed=1; low=2). Postnatal echocardiographic data was reviewed where available to confirm diagnosis.

Postnatal neurodevelopmental outcomes were assessed at Boston Children’s Hospital between 14.8 to 39.3 months of age using either the third or fourth (Bayley) edition of the Bayley Scales of Infant and Toddler Development^42^, reflecting the transition in the institutional assessment clinical practice. Analyses included gross motor, fine motor, expressive language, receptive language, and cognition scaled scores (mean=10, SD=3), with higher scores indicating better developmental performance and scores <7 indicative of developmental delay.^42^ Because Bayley edition had significant effects on developmental scores and age at assessment demonstrated smaller but non-negligible effects (SI-Table1), both variables were included as covariates in analyses involving neurodevelopmental outcomes.

### 2.2. Fetal MRI acquisition and processing

MRI were performed on 3T Siemens scanners (MAGNETOM Skyra and Prisma; Siemens Healthineers, Germany). Imaging consisted of repeated multi-planar T2-weighted half-Fourier single-shot turbo spin echo (HASTE) acquisitions optimized for fetal neuroimaging.

Sequence parameters included repetition time (TR)=1400-1600ms, echo time (TE)=99-132ms, 1 mm in-plane resolution, 2-3.5 mm slice thickness, and a variable field of view adapted to fetal and maternal anatomy. For each subject, HASTE stacks were acquired in axial, coronal, and sagittal orientations, with at least three repeated acquisitions per orientation (∼30 minutes total acquisition time).

Image processing was performed using a dedicated fetal MRI reconstruction pipeline. Processing steps included fetal brain extraction^43^, N4 bias field correction^44^, and slice-to-volume reconstruction using NeSVoR.^45^ These procedures generated motion-corrected super-resolved 3D T2-weighted volumes at 0.5 mm isotropic resolution. Additional details regarding HASTE stack selection and reconstruction quality are provided in SI-Fig1 and SI-Fig4.

SP segmentation was performed semi-automatically using a previously described deep-learning-based framework.^32^ Initial SP labels generated by the network were visually inspected and manually corrected using the underlying T2-weighted images to address segmentation inaccuracies. Although all segmentations were corrected by a single trained reader, inter-reader and intra-reader variability analyses are provided in SI-Fig5 for reference. In addition to the SP, the cortical plate (CP) and an “Other” tissue class, comprising the intermediate zone (IZ) and remaining supratentorial tissues, were also delineated to provide anatomical context and facilitate comparison with prior fetal neuroimaging studies.

Outer SP surfaces, corresponding to the CP/SP boundary, were generated using the ChRIS pl-fetal-surface-extract plugin^46^, which implements a CIVET-like marching cubes algorithm^47^, followed by Taubin smoothing.^48^ Inner SP surfaces, corresponding to the SP/IZ boundary, were derived by inward deformation of the outer surface along subject-specific radial distance maps generated from the inner SP boundary.^49^ This approach preserved vertex-wise correspondence between inner and outer SP surfaces. Following extraction, surfaces were resampled to standardized meshes consisting of 81,920 triangles.

Surface quality control was performed both visually and quantitatively. Quantitative metrics included smoothness error, defined as the mean curvature deviation between each vertex and its neighboring vertices, and boundary distance error, defined as the Euclidean distance between each surface vertex and the nearest boundary voxel. These metrics were used to identify subjects requiring closer visual inspection. No subjects were excluded on the basis of surface quality (SI-Fig6).

### 2.3. Regional parcellation

For regional analyses, 21 bilateral cortical regions were delineated using a fetal-specific adaptation of the Desikan-Killiany atlas^50^, as previously described^9,32^ (SI-Fig7A). Regional parcellations were manually defined on a 29w surface template^51^ and subsequently propagated to individual subject surfaces using a two-dimensional spherical surface registration approach.^52^ This methodology has been successfully applied in several prior fetal surface-based studies.^10,32,53^ Visual quality control demonstrated globally consistent regional delineation across subjects and confirmed the absence of gross registration failures (SI-Fig7B).

Following regional quality control, four bilateral regions were excluded from subsequent analyses: (i) the orbitofrontal cortex, due to frequent frontoventral blurring on T2-weighted MRI; and (ii) the cingulate gyrus, insula, and parahippocampal gyrus, due to surface extraction artifacts arising from abrupt SP label termination that resulted in local over- or under-estimation of surface geometry. Consequently, 17 bilateral cortical regions were retained for regional analyses.

For lobar analyses, regional measures were additionally combined into frontal, temporal, parietal, and occipital lobe summaries according to the assignments shown in SI-Fig7B.

### 2.4. Quantification of whole-brain and regional SP volumes and thickness

Whole-brain CP, SP, and “Other” tissue volumes were calculated as the number of voxels within each tissue label multiplied by voxel volume in native space. Global supratentorial brain volume (“brain volume”) was additionally computed as the sum of all tissue classes. Whole-brain SP thickness was defined as the Euclidean distance between corresponding vertices on the inner and outer SP surfaces and summarized as the median value across both hemispheres, excluding the cingular pole.

Regional measures were computed for the 17 bilateral cortical regions described above. To derive regional volumetric measures, surface-based parcellations were projected into volumetric space using a ribbon-constrained approach restricted to the SP label and adapted from Connectome Workbench (v2.0.1). Regional SP volume was then calculated as the number of voxels assigned to each region multiplied by voxel volume. Regional SP thickness was defined as the median vertex-wise thickness value within each region. This SP measurement pipeline has previously been successfully applied to characterize typical SP development in fetal brain MRI^32^.

Because regional SP thickness may vary systematically with local surface geometry, we additionally quantified SP depth to account for folding-related effects. SP depth, reflecting the amplitude of cortical folding along the outer SP surface, was adapted from Makropoulos et al.^54^ and implemented using MIRTK (v2.0). Specifically, SP depth was calculated as the displacement along surface normals between the native outer SP surface and an inflated representation of the same surface. Depth values were rescaled such that the minimum hemispheric depth value was zero for each subject, excluding the cingular pole. Regional SP depth was summarized as the median vertex-wise depth value within each region.

For lobar analyses, regional measures were aggregated within each lobe by summing regional SP volumes and taking the median of regional SP thickness and depth values according to the regional assignments shown in SI-Fig7B. Raw whole-brain, lobar, and regional SP volume and thickness measures are visualized in SI-Fig8.

### 2.5. Statistical analysis

All SP volumetric and thickness measures were natural log-transformed prior to analysis to improve normality and stabilize variance. Where applicable, p-values were adjusted for multiple comparisons using the Benjamini-Hochberg false discovery rate (FDR) procedure, with adjusted values reported as q-values. Details of FDR correction are provided in the corresponding Results sections.

#### 2.5.1 Group-level and hierarchical modelling

Whole-brain SP measures were first modeled using linear regression with group (CHD vs typically developing controls, TD), GA, and sex as fixed effects. At hemispheric, lobar, and regional levels, linear mixed-effects models with subject-specific random intercepts were fitted to account for repeated measures within subjects.

##### SP thickness and SP depth adjustment

For SP thickness analyses, SP depth was included as an additional covariate to account for geometric effects of cortical folding.^32^ In short, SP depth was decomposed into *between-subject* (mean sulcal depth per subject) and *within-subject* (deviation from subject mean) components. Their associations with SP thickness were evaluated using mixed-effects models including interactions with hemisphere, lobe, and region, separately.

The within-subject SP depth component showed significant interactions with lobes (F=494.25, p<0.001, η²=0.10) and regions (F=379.25, p<0.001, η²=0.08), but not hemispheres (F=0.93, p=0.335, η²<0.01). Between-subject SP depth effects were not significant (p>0.411). Accordingly, only the within-subject component was retained in final sub-whole-brain models, including hemispheres for consistency (SI-Table 3).

##### Global brain size adjustment

To account for global brain volume effects, extended models were fitted at all modelling levels. At the whole-brain SP volume and thickness level, residual brain volume adjusted for GA and sex was included as a covariate. At hemispheric, lobar, and regional levels, SP volume models additionally included whole-brain SP volume, and SP thickness models included median whole-brain SP thickness (both adjusted for GA and sex). These models were used to isolate regional effects beyond global scaling of brain size or overall SP measures. Estimated marginal means were extracted from fitted models and back-transformed from the log scale to yield geometric mean estimates of SP volume or thickness in CHD and TD groups.

##### Gestational age modelling

Given the potential non-linear relationship between SP metrics and GA, we compared models specifying GA as either a linear term or using restricted splines (up to 4^th^ order). Final model specification was selected based on Akaike Information Criterion (AIC), with Bayesian Information Criterion (BIC) used as a secondary criterion when model selection was ambiguous.

#### 2.5.2 Developmental deviation analysis (TD-referenced framework)

To quantify deviations from typical SP developmental trajectories, we constructed a TD-referenced normative modelling framework. Expected developmental trajectories were estimated using TD subjects only.

At the whole-brain level, log-transformed SP measures were modelled as a function of GA and sex, with residual brain volume included in a second step. At sub-whole-brain levels, mixed-effects models similarly included residualized whole-brain SP volume or median SP thickness to account for global effects.

The TD-trained model was then applied to all subjects to generate predicted values. Individual deviations were computed in log-space as observed minus predicted values, representing deviation from expected developmental trajectories. To enable standardized comparisons, residuals were further converted into Z-scores using the mean and standard deviation of the TD residual distribution, with negative and positive values indicating lower- and higher-than-expected SP measures, respectively. Group differences in deviations were assessed using Wilcoxon rank-sum tests applied to both signed residuals and absolute Z-scores to capture differences in direction and magnitude. The proportion of individuals exhibiting extreme deviations (|Z|>1.96) was compared using Fisher’s exact test.

#### 2.5.3 SP deviations in CHD and measures of fetal cardiac hemodynamics

We next examined associations between SP developmental deviations and fetal cardiac physiology, including cerebroplacental hemodynamics and substrate delivery metrics. Analyses focused on TD-referenced SP deviation measures derived from the normative modelling framework.

At the whole-brain level, linear regression models were fitted separately for SP volume and SP thickness deviations. First, substrate delivery score and CPR were modeled independently (with Echo-MRI delay as a covariate), then jointly. Additional models evaluated associations after further adjustment for residual whole-brain volume to isolate effects beyond global brain-size variation.

At hemispheric, lobar, and regional levels, linear mixed-effects models were used to account for repeated measures within subjects. These models included subject-specific random intercepts and fixed effects for region, cardiac variables, and their interactions to assess spatial heterogeneity of associations across the cortex. Given the limited sample size for subgroup analyses and the exploratory nature of these analyses, findings were interpreted cautiously.

#### 2.5.4 SP deviations in CHD and neurodevelopmental outcomes

Associations between prenatal SP developmental deviations and postnatal neurodevelopmental outcomes were assessed in CHD subjects with available Bayley data. Analyses were performed separately for whole-brain and regional SP deviation measures.

At the whole-brain level, linear regression models were fitted with Bayley scaled scores as dependent variables and SP deviation measures as predictors. Base models included Bayley edition and age at assessment as covariates. Because neurodevelopment in CHD is also shaped by perioperative, clinical, and environmental factors not captured by fetal MRI, we additionally included GA at birth, duration of neonatal hospitalization, history of neonatal seizures, and primary parent’s education level as covariates.^19^

At the regional level, separate linear regression models were fitted for each cortical region and developmental domain. Predictor variables included regional TD-referenced SP volume or thickness deviations (GA, sex, and whole-brain SP measure-adjusted). Associations were evaluated across five developmental domains: gross motor, fine motor, expressive communication, receptive communication, and cognition.

Given the large number of comparisons and limited longitudinal sample size, these analyses were considered exploratory. P-values were corrected for multiple comparisons using the Benjamini-Hochberg FDR procedure within each outcome domain and are denoted as q-values.

## 3. Results

### 3.1. Cohort characteristics

The final cohort included 76 singleton fetuses with CHD (50 male; median GA at MRI=28.8wGA [range: 21.1–31.9]) and 62 TD singleton fetuses (33 male; median GA at MRI= 27.5wGA [range: 22.0–31.4]). Groups did not differ significantly in GA at MRI or sex distribution. As Doppler assessment was performed significantly earlier in the TD than in the CHD group (median GA et Echo: 22.1 vs 27.1wGA, respectively, p<0.001), CPR values were standardized for GA. The ECHO-MRI interval was also longer in TD than CHD fetuses (median 4.9 vs 1.2 weeks; p < 0.001). CHD fetuses showed a higher prevalence of abnormal CPR (Z<-1: 17% vs 4%, p=0.037) (Table 1). Examination of the individual CPR components showed a trend toward higher UA PI in CHD fetuses, whereas MCA PI did not differ significantly between groups (SI-Table2).

**Table 1.** Select subject characteristics. Differences between TD and CHD groups for continuous variables were assessed using t-tests with Welch’s correction for unequal variances (t: t-statistic, dof: degrees of freedom, p: p-value, D: Cohen’s D). For categorical variables, differences were evaluated using Chi-squared (Χ²) tests (Χ²: Χ²-statistic, V: Cramer’s V). For neurodevelopmental measures, differences are evaluated using linear models including Bayley edition and age at assessment as covariates (t: t-statistic, dof: degrees of freedom, q: p-value, FDR-corrected, D: Cohen’s D). N: number of subjects. See SI-Table1 for additional cohort information.

|  | TD (N=62) | CHD (N=76) |  |
| --- | --- | --- | --- |
| **Fetal characteristics:** |  |  |  |
| GA at scan (weeks): median [range] | 27.5 [22.0-31.4] | 28.8 [21.1-31.9] | t=-1.76, dof=136.00,<br>p=0.081, D=0.30 |
| Males/Females: count (%) | 33 (53) /29 (47) | 50 (66) /26 (34) | $\chi^2=1.75$, dof=1.00,<br>p=0.185, V=0.11 |
| GA at Echo (weeks): median [range] (N) | 22.1 [18.0-31.6] (60) | 27.1 [20.4-36.4] (74) | **t=-9.18, dof=131.55,**<br>**p<0.000, D=1.55** |
| Echo-MRI delay (weeks): median [range] (N) | 4.9 [0.9-11.0] (60) | 1.2 [0.0-6.0] (74) | **t=10.51, dof=95.29,**<br>**p<0.000, D=1.91** |
| Cerebroplacental ratio (Z): median [range] | -0.5 [-1.3-0.1] (55) | -0.6 [-1.7-0.9] (70) | t=0.42, dof=112.41,<br>p=0.678, D=0.07 |
| CPR abnormality (Z<-1): count (%) | 2 (4) | 12 (17) | **$\chi^2=4.37$, dof=1.00,**<br>**p=0.037, V=0.19** |
| **Cardiac characteristics:** |  |  |  |
| Diagnosis: count (%)<sup>a</sup> |  |  |  |
| HLHS | - | 12 (16) |  |
| TGA | - | 18 (24) |  |
| TOF | - | 14 (18) |  |
| FAS | - | 7 (9) |  |
| Other two ventricle | - | 16 (21) |  |
| Other single ventricle | - | 9 (12) |  |
| Single ventricle physiology: count (%) | - | 22 (29) |  |
| Substrate delivery score: count (%) |  |  |  |
| 0 |  | 25 (33) |  |
| 1 |  | 8 (10) |  |
| 2 |  | 27 (36) |  |
| 3 |  | 16 (21) |  |
| **Neurodevelopmental outcomes:** |  |  |  |
| Bayley at 18-24m: count (%) | 37 (60) | 46 (61) | - |
| Bayley-III (%) | 20 (54) | 30 (65) | $\chi^2=0.65$, dof=1.00,<br>p=0.419, V=0.09 |
| Bayley-IV (%) | 17 (46) | 16 (35) |  |
| Age at Bayley (months): median [range] | 20.0 [19.4-39.3] (37) | 21.7 [14.8-37.6] (46) | t=0.89, dof=71.82,<br>p=0.376, D=0.20 |
| Scaled scores: median [range] |  |  |  |
| Gross motor | 9 [6-17] | 9 [3-12] | **t= 3.28, dof=79,**<br>**q=0.008, D=0.74** |
| Fine motor | 11 [7-17] | 10 [5-15] | t=1.59, dof=79,<br>q=0.122, D=0.36 |
| Expressive language | 10 [4-19] | 9 [1-16] | t=1.57, dof=79,<br>q=0.122, D=0.35 |
| *Receptive language* | 12 [1-19] | 10 [4-15] | $t=1.84$, $dof=79$,<br>$q=0.117$, $D=0.41$ |
| *Cognitive* | 11 [7-18] | 10 [1-14] | $t=2.60$, $dof=79$,<br>$q=0.028$, $D=0.59$ |

<sup>a</sup>HLHS: hypoplastic left heart syndrome; TGA: d-transposition of the great arteries; FAS: fetal aortic stenosis; TOF: tetralogy of Fallot.

Of the 76 CHD fetuses, 7 (11%) died shortly after birth. Among the surviving participants, neurodevelopmental outcomes were available for 46 CHD subjects (61% of the full cohort) and 37 TD subjects (60%). One additional subject has not yet completed follow-up but remains within the eligible assessment age window; follow-up assessments were unavailable for the remaining participants due to loss to follow-up, withdrawal, or inadequate data quality. Compared with TD peers, CHD subjects demonstrated significantly lower gross motor and cognitive scaled scores (Table 1, SI-Fig3C).

Expanded maternal, fetal, and postnatal characteristics are provided in SI-Table2 and SI-Fig3.

### 3.2. Group-level differences in SP volume

Consistent with expected fetal developmental trajectories, both TD and CHD groups showed gestational age (GA)-related increases in total brain volume and in individual tissue compartments, including cortical plate (CP), subplate (SP), and “Other” tissue volumes (SI-Fig9).

Whole-brain SP volume increased strongly with GA (F=713.32, p<0.001, η²=0.91) and differed by sex (F=15.98, p<0.001, η²=0.12). CHD fetuses showed lower SP volumes overall, although the group effect did not reach significance after accounting for GA using second-order splines (F=2.82, p=0.096, η²=0.02; SI-Table4A). Regional analyses demonstrated spatially heterogeneous reductions in SP volume in CHD, with significant reductions observed in the left hemisphere (p=0.024, Cohen’s D=-1.69), particularly within parietal-occipital regions and right temporal cortex (SI-Table4B; Fig1A).

**Fig 1.**
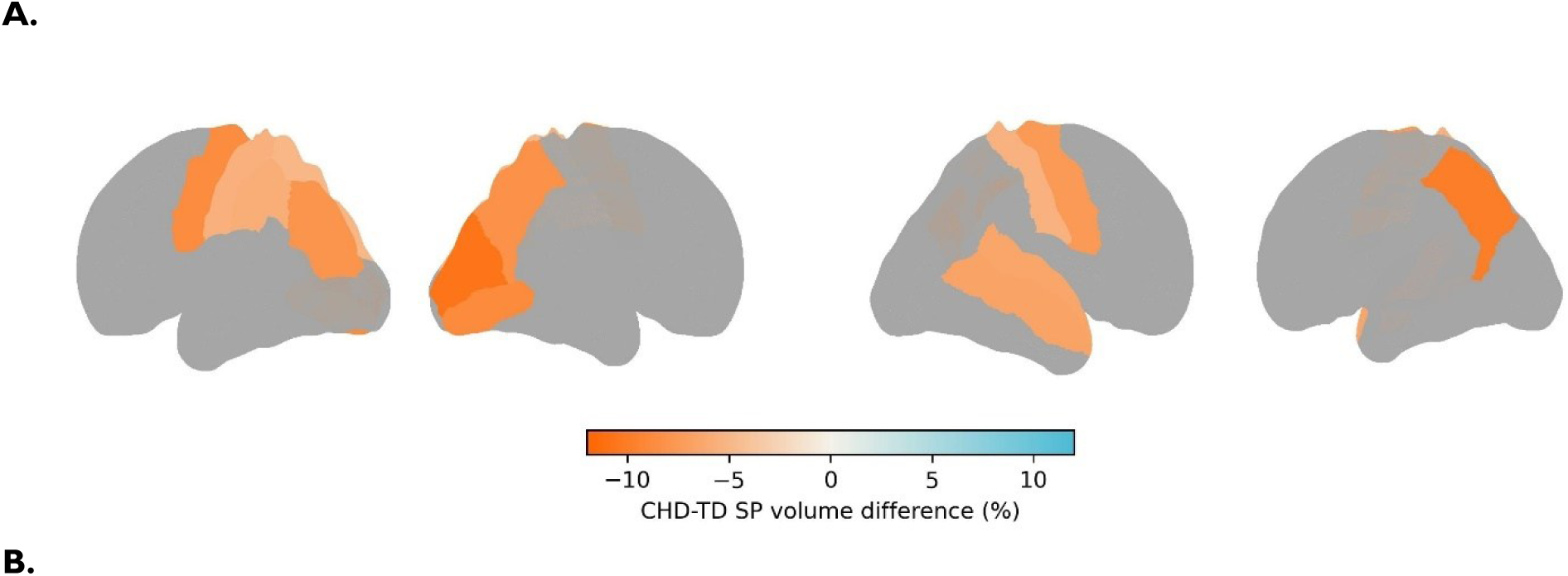
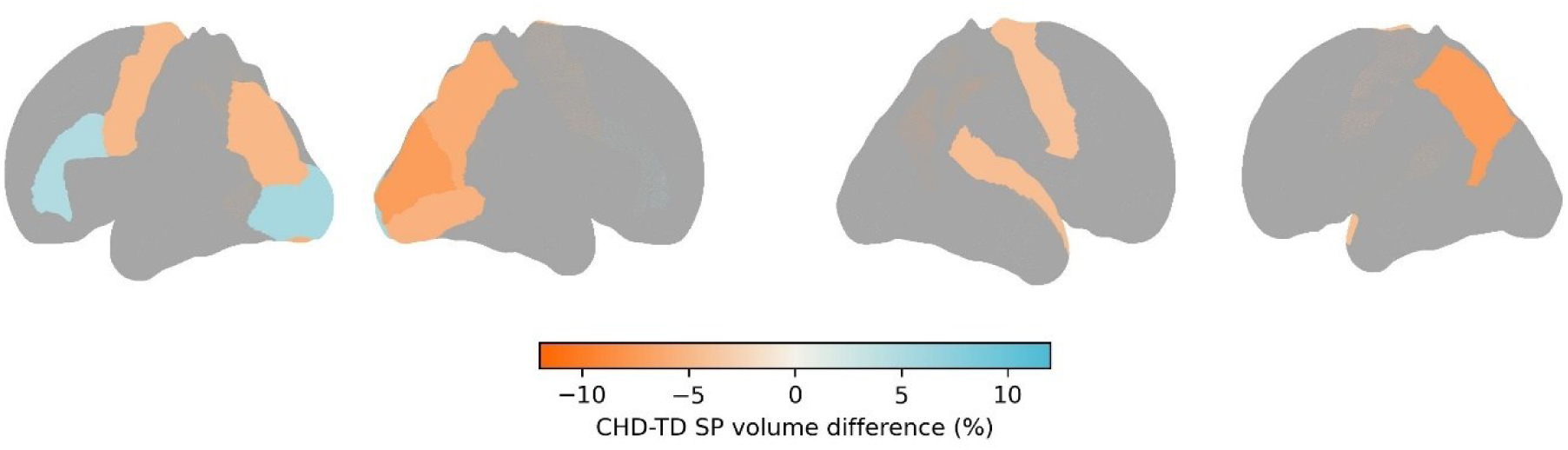
Regional differences in subplate volume. **A.** Surface maps showing regional CHD-TD differences in SP volume identified in ANCOVA analyses (FDR-corrected; predicted at mean GA=27.8 weeks, male sex). Warm colors indicate lower SP volume in CHD relative to TD. **B.** Regional SP volume differences additionally adjusted for residual whole-brain SP volume (adjusted for GA and sex) (predicted at mean SP volume=34.26 cm^3^).

Given the significant global reduction of brain volumes in CHD vs TD fetuses (GA and sex adjusted; Welch’s t=-3.52, p=0.001, Cohen’s D=-0.55), we next examined whether SP alterations exceeded overall brain-size differences. After additionally accounting for residual whole-brain volume, the group effect on whole-brain SP volume was no longer significant (F=0.48, p=0.492, η²<0.01; SI-Table4C), indicating that much of the observed SP reduction aligned with global reductions in brain volume. However, regional analyses adjusted for whole-brain SP volume continued to demonstrate localized deviations, including relatively smaller bilateral precentral gyri, precuneus, left inferior parietal cortex, cuneus, lingual gyrus, and right superior temporal cortex (SI-Table4D; Fig1B). These findings suggest that regional SP alterations in CHD are superimposed on a dominant global reduction in brain size.

### 3.3. Group-level differences in SP thickness

Whole-brain SP thickness increased with GA (F=212.64, p<0.001, η²=0.87) and differed by sex (F=11.89, p<0.001, η²=0.08), but not group (F=0.23, p=0.331, η²<0.01; SI-Table5A). Whole-brain SP thickness was also associated with global brain volume (F=29.76, p<0.001, η²=0.19), further reducing independent group effects after adjustment (SI-Table5C).

Regional analyses revealed spatially heterogeneous reductions in SP thickness in CHD, predominantly affecting the left hemisphere (p=0.041, Cohen’s D=-0.69), especially occipital and parietal regions (SI-Table5B; Fig2A). After accounting for whole-brain SP thickness, localized reductions remained primarily within medial posterior cortex, superior parietal cortex, and middle temporal gyrus (SI-Table5D; Fig2B). Overall, the pattern was consistent with relatively greater involvement of posterior cortex and temporal regions in CHD.

**Fig 2.**
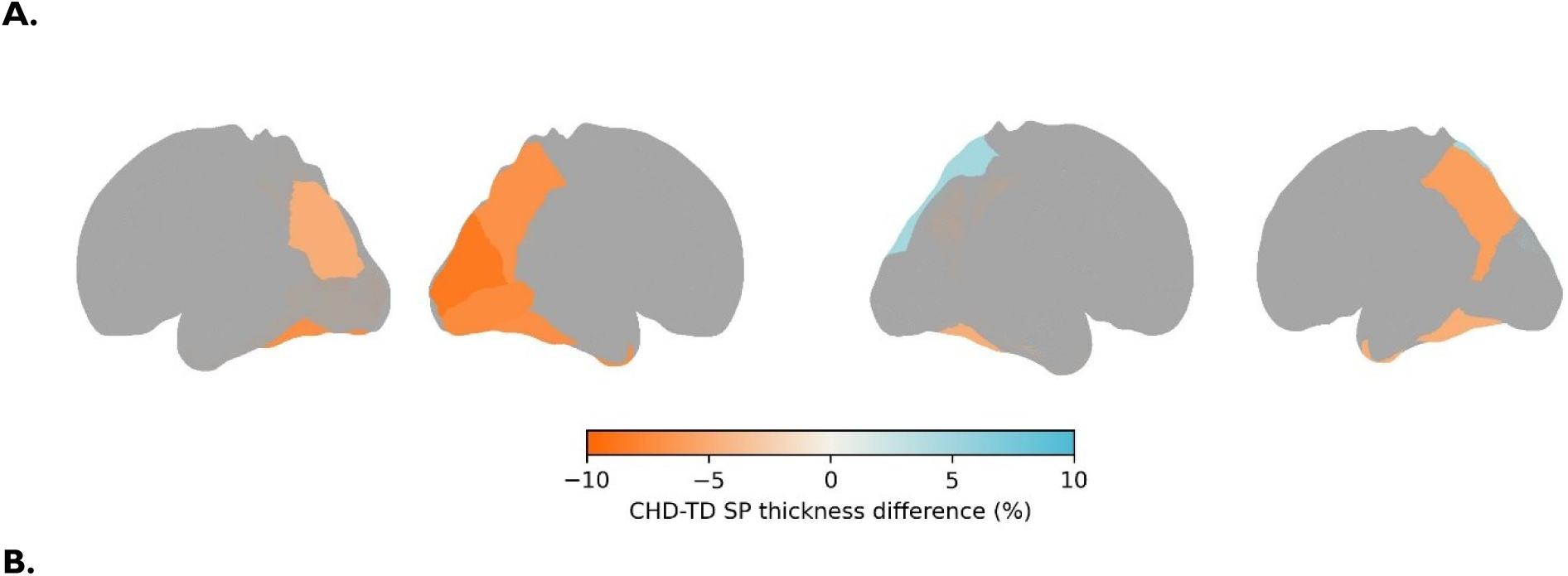
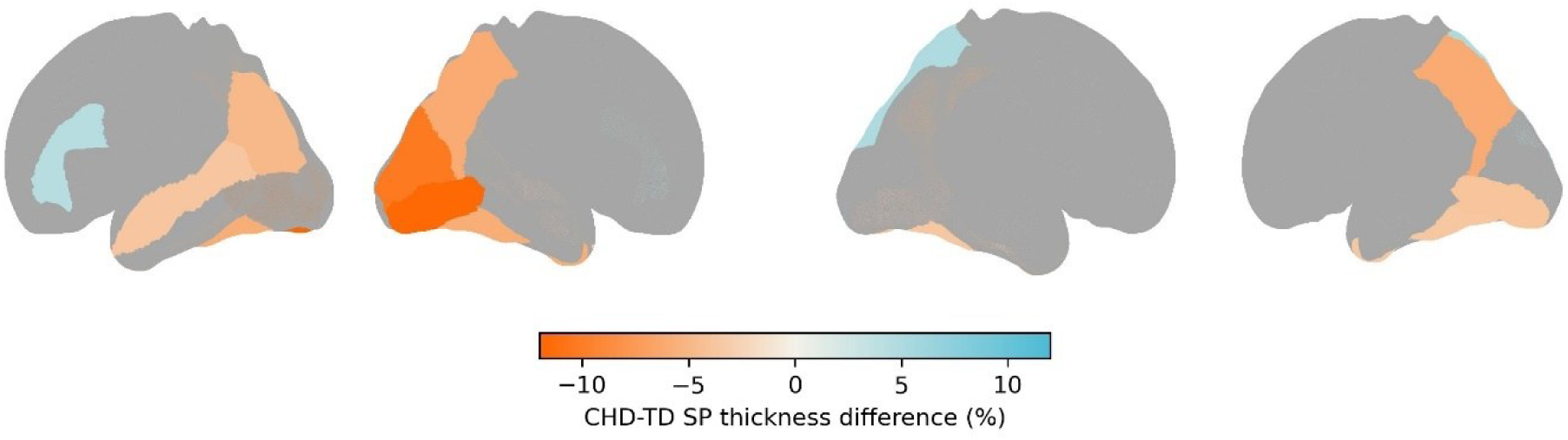
Regional differences in subplate thickness. **A.** Surface maps showing regional CHD-TD differences in SP thickness identified in ANCOVA analyses (FDR-corrected; predicted at mean GA=27.8 weeks, male sex, SP depth=6.08mm). Warm colors indicate lower SP thickness in CHD relative to TD. **B.** Regional SP thickness differences additionally adjusted for residual whole-brain SP thickness (adjusted for GA and sex) (predicted at mean SP thickness=3.33mm, SP depth=6.08mm).

### 3.4. Normative modelling of developmental deviations

#### 3.4.1 Whole-brain deviations

Relative to normative regional brain developmental trajectories, CHD fetuses showed significantly greater absolute deviations in whole-brain SP volume (Wilcoxon W=3183, p<0.001, rank-biserial correlation=0.35), with systematic reductions relative to GA- and sex-expected values (W=1673, p=0.003, rank-biserial correlation=-0.29; SI-Fig10A). These reductions corresponded to an average SP volume reduction of 5.7% relative to expected values. Extreme deviations (|Z|>1.96) were observed in 16% of CHD fetuses compared with 5% of TD fetuses, although this difference did not reach significance (Fisher’s exact test, odds ratio=0.27, p=0.054).

Within CHD, HLHS/TGA fetuses showed larger SP volume deviations than Other-CHD fetuses, with only the HLHS/TGA subgroup differing significantly from TD controls (t=-4.21, p<0.001; SI-Fig10A). However, after accounting for residual whole-brain volume, group differences in SP volume deviations were attenuated, with no remaining differences in deviation magnitude (Wilcoxon Rank Sum test, W=2753, p=0.090, rank-biserial correlation=0.17), directional bias (Wilcoxon Rank Sum test, W=2359, p=0.992, rank-biserial correlation<0.01), or prevalence of extreme deviations (12% in CHD vs 6% in TD, Fisher’s exact test, odds ratio=0.52, p=0.383). These findings again indicate SP volume reductions are proportional to global reductions in brain size.

For whole-brain SP thickness, deviations did not differ significantly between CHD and TD groups (SI-Fig10B). Although extreme SP thickness deviations were more frequent in CHD (11% vs 2%, Fisher’s exact test, odds ratio=0.14, p=0.041), these differences were also attenuated after accounting for residual brain volume. Together, these findings suggest that whole-brain SP alterations in CHD are proportional to global reductions in brain growth rather than disproportionate abnormalities in SP thickness.

#### 3.4.2 Regional deviations

Regional normative deviations demonstrated marked spatial heterogeneity across the cortex (Fig3). After adjustment for GA, sex, and whole-brain SP volume, CHD fetuses showed regionally selective SP volume deviations relative to TD expectations. Positive deviations were observed predominantly in right frontal (Z=0.12±1.101, T=2.64, Cohen’s D=0.10, p=0.008) and occipital lobes (Z=0.20±1.047, T=3.96, Cohen’s D=0.19, p<0.001), including lateral occipital cortex, cuneus, and superior, middle, and inferior frontal gyri. Negative deviations were concentrated within left parietal lobe (Z=-0.13±1.008, T=-2.72, Cohen’s D=0.10, p=0.007) and occipital regions, including inferior parietal cortex, precuneus, cuneus, and lingual gyrus (SI-Table 6A; Fig 3A).

**Fig 3.**
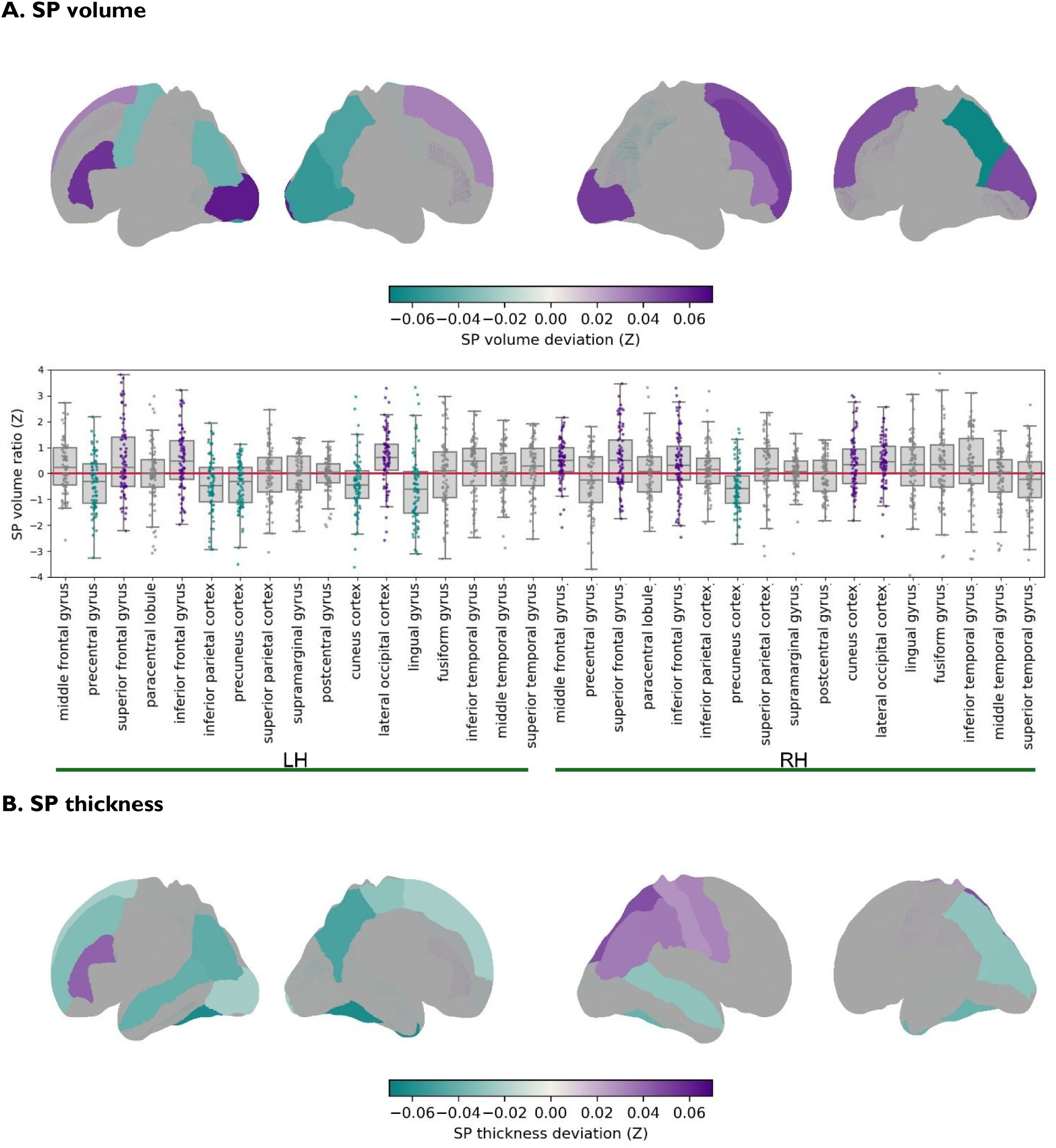
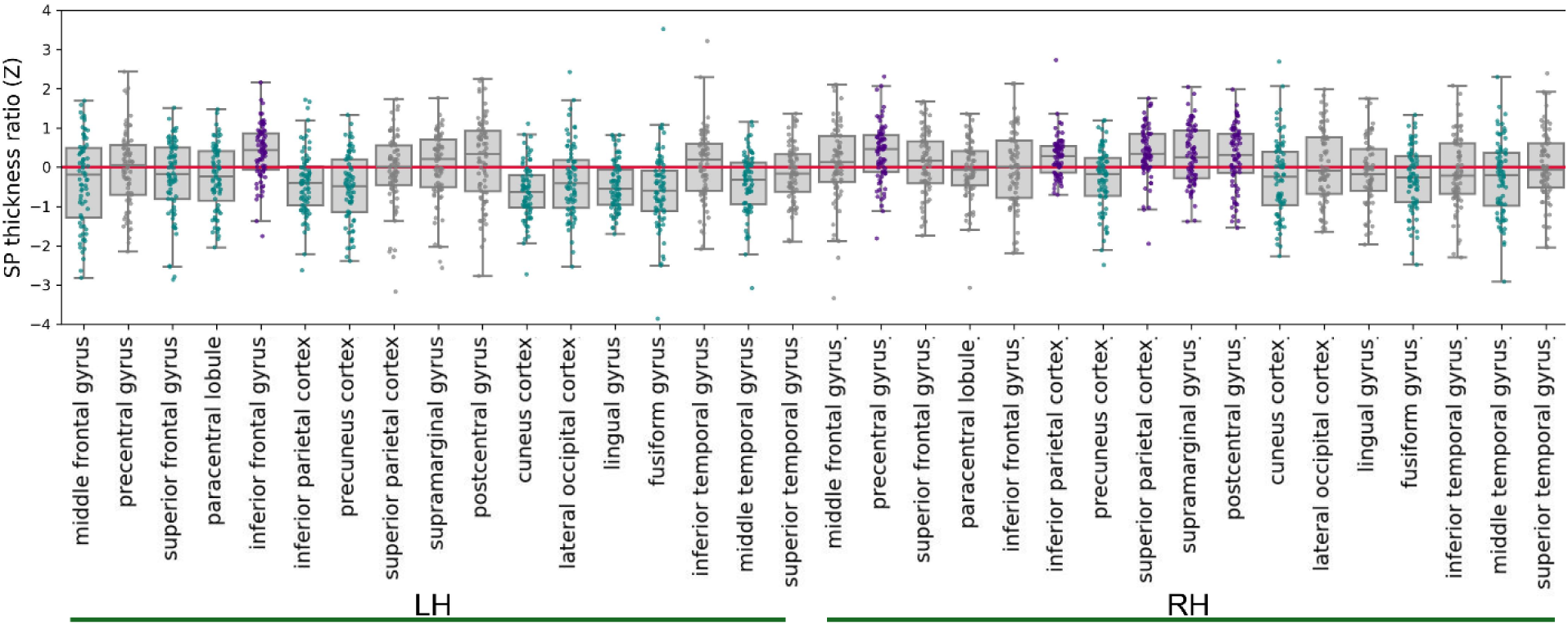
Regional normative subplate deviations in CHD. **A**. Regional SP volume deviations relative to TD normative trajectories. Surface maps show mean regional Z-scores across CHD subjects. Boxplots summarize group distributions, and scatter points represent individual subjects. Only regions showing significant deviations from zero (one-sample t-tests, FDR-corrected) are displayed in color, with negative deviations shown in turquoise and positive deviations in purple. **B.** Regional SP thickness deviations displayed using the same format.

SP thickness deviations showed a partially overlapping but distinct spatial distribution (SI-Table6B; Fig3B). Negative deviations were most prominent within left hemisphere (Z= −0.013±0.970, T=-6.86, Cohen’s D=0.14, p<0.001) across parietal, occipital, and temporal lobes, including precuneus, fusiform gyrus, and middle temporal regions. Significant positive deviations were observed across regions within right parietal lobe (Z=0.10±0.870, T=3.40, Cohen’s D=0.13, p=0.001) (SI-Table6B; Fig3B).

Substantial inter-individual variability was observed across subjects. Extreme regional deviations (|Z|>1.96) in SP volume were enriched in CHD, with 73% of cortical regions demonstrating odds ratios >1 for extreme deviations relative to TD controls (binomial test p=0.009). Significant enrichment after FDR correction was observed primarily within left lingual, fusiform, and superior frontal gyri (SI-Fig11A). Enrichment of extreme SP thickness deviations was less widespread (58% of regions showing OR>1; p=0.392), although elevated odds ratios were observed across bilateral parietal and temporal regions (SI-Fig11B). Together, these findings suggest that regional SP deviations in CHD are spatially heterogeneous across individuals, with variable patterns of atypical cortical development across the cohort, consistent with the underlying heterogeneity of CHD physiology.

### 3.5. Associations between SP deviations in CHD and fetal hemodynamics

We next examined associations between SP developmental deviations in CHD, cerebral substrate delivery score, and CPR Z-scores. Neither measure was associated with SP volume deviations, although CPR showed a trend-level association after adjustment for brain size (F=3.82, p=0.055, η²=0.06 β=-0.55). For GA- and sex-referenced deviations SP thickness deviations, CPR showed a trend-level negative association (F=3.61, p=0.062, η²=0.05, β=-0.52), which strengthened after adjustment for substrate delivery score (F=4.58, p=0.036, η²=0.07, β= −0.59). After additionally accounting for brain size, CPR remained negatively associated with SP thickness deviations (F=6.06, p=0.017, η²=0.08, β=-0.65), with a stronger association in the joint model including substrate delivery score (F=7.07, p=0.010, η²=0.10, β=-0.59). Substrate delivery score alone did not reach significance in any model (SI-Table7).

Substrate delivery score (substrate delivery score*region; F=1.52, p=0.030, η²=0.03) was suggested to have regionally heterogenous effects on SP development. However, no regional effects survived correction for multiple comparisons, and these analyses should therefore be considered hypothesis-generating.

Together, these analyses suggest that CPR may be associated with SP developmental deviations in CHD, most consistently for SP thickness and after accounting for cerebral substrate delivery. Given the modest effect sizes, these findings remain exploratory.

### 3.6. Associations between SP deviations in CHD and neurodevelopmental outcomes

Associations between prenatal SP deviations in CHD subjects and two-year neurodevelopmental outcomes were examined using Bayley scaled scores (while accounting for test edition and age at assessment as well as GA at birth, duration of neonatal hospitalization, history of neonatal seizures, and primary parent’s education level as covariates). At the whole-brain level, SP volume deviations (GA and sex TD-referenced) showed association with receptive language scores (F=4.93, p=0.033, η²=0.12) but not cognitive, fine or gross motor scores (SI-Fig12A). Fully corrected SP volume deviations (i.e. GA, sex, whole-brain volume TD-referenced) showed a significant association with fine motor scores (F=8.23, p=0.007, η²=0.19) and a nominal relationship with cognitive scores (F=3.95, p=0.055, η²=0.10) but not gross motor, expressive or receptive language scaled scores (SI-Fig12A). No significant associations were observed for whole-brain SP thickness deviations.

Regionally, SP volume deviations showed nominally significant associations (uncorrected p<0.05) with receptive communication and cognitive scores in temporal regions, expressive communication, cognitive, and gross motor outcomes in frontal regions, and expressive/receptive communication in lateral occipital cortex. SP thickness deviations clustered in parietal regions, relating to cognitive, language, and fine motor outcomes (SI-Table8, SI-Fig12). Only right precuneus SP thickness deviations survived FDR correction, remaining associated with receptive (β=-11.87, q=0.017) and expressive (β=-14.50, q=0.039) communication scores; all other regional associations should be interpreted as exploratory.

## 4. Discussion

This study provides evidence that SP development in CHD is altered in a spatially heterogeneous and individually variable manner during the late second and early third trimester, superimposed on a dominant context of global brain size reduction. Using complementary volumetric and surface-based analyses together with TD-referenced normative developmental modelling, we demonstrate that while much of the observed SP reduction scales with overall brain size, regional abnormalities persist beyond global adjustment, particularly in posterior parietal, occipital, and temporal cortices, and vary substantially between individuals. These findings raise the possibility that altered fetal physiology may interact with regionally distinct maturational programs during a critical developmental window. Associations between SP deviations, fetal hemodynamic indices, and two-year neurodevelopmental outcomes further highlight potential links between prenatal SP development and later neurodevelopment, although these analyses remain exploratory.

### 4.1. Global reductions in brain size as the dominant feature of SP alterations

The reduction in whole-brain SP volume is consistent with previous fetal MRI studies reporting smaller brains and altered cortical and SP volumes in CHD.^4,8^ Importantly, attenuation of the SP volume difference after adjustment for global brain volume indicates that much of the apparent global SP reduction scales with overall cerebral size. The absence of a corresponding whole-brain SP thickness difference further supports this interpretation. Thus, our findings do not support a generalized disproportionate reduction of the SP in CHD but rather suggest that global SP changes occur largely in the context of reduced overall brain growth previously observed across tissue compartments.^6–8^

### 4.2. Regional SP alterations: spatial structure and hemispheric asymmetry

Regional analyses provided evidence for deviations beyond the global brain reductions in CHD. After accounting for global SP measures, SP volume and thickness differences were most consistently observed in posterior parietal, occipital, and temporal regions, with additional involvement of the precentral gyrus, and were particularly pronounced in the left hemisphere for SP thickness. These regions undergo substantial SP expansion during the second and third trimesters^32,55^, and may therefore be sensitive to altered fetal conditions during a period of increasing metabolic demand.

The spatial pattern of regional deviations may relate to known posterior-to-anterior and medial-to-lateral maturational hierarchies, with association cortices showing more protracted and heterogeneous developmental trajectories relative to primary motor and sensory regions.^56^ The relatively increased SP volume in occipital regions after global adjustment may reflect relative preservation of occipital SP despite global reductions. We have previously described similar hierarchical patterns in SP development.^32^ Intrinsic regional differences in cellular composition, molecular identity, glial distribution, and connectivity may further shape local developmental trajectories and vulnerability to altered fetal conditions^31,57^ although their specific contribution in CHD remains unknown.

Prenatal hemispheric asymmetries have been documented across cortical growth, gyrification, structural connectivity, and SP morphology.^9,32,58,59^ The partial left-hemisphere predominance observed here raises the possibility that altered fetal physiology interacts with ongoing hemispheric differentiation. However, the cross-sectional design precludes determining whether these asymmetries reflect progressive developmental differences, maturational timing, or sampling variation.

The spatial correspondence of these prenatal findings with regions implicated in postnatal CHD neuroimaging provides a useful reference. Reduced cortical volumes have been reported in superior frontal, middle frontal, and parietal regions, while altered cortical thickness has been observed in distributed regions including parietal and cingulate cortices.^60,61^ However, because the SP undergoes perinatal resolution, direct structural comparison with postnatal MRI is limited. It therefore remains unclear whether prenatal SP alterations contribute to later cortical abnormalities or instead reflect shared regional vulnerability across developmental stages.

### 4.3. Normative modelling reveals individualized patterns of SP development

A further contribution of the present study is the application of TD-referenced normative modelling to characterize inter-individual variability in SP development. This is particularly relevant in CHD, where cardiac diagnoses encompass heterogeneous circulatory phenotypes, substrate delivery profiles, and genetic backgrounds. At the whole-brain level, fetuses with CHD showed SP volume reduction (∼5.7% relative to GA- and sex-expected TD values), which was substantially attenuated after accounting for global brain volume, consistent with group-level analyses. CHD fetuses nevertheless showed a higher frequency of extreme deviations, particularly for SP thickness, and enrichment of extreme regional volume deviations across most regions, consistent with prior fetal MRI studies demonstrating substantial heterogeneity of brain alterations across CHD diagnoses and individuals.^8,14,62,63^

Regional SP deviations were bidirectional and spatially structured after adjustment for GA, sex, and whole-brain SP measures. Volume deviations were primarily negative in medial posterior regions including the precuneus, inferior parietal cortex, and lingual gyrus, with positive deviations in lateral occipital and right frontal cortex. Thickness showed a distinct, predominantly left-lateralized pattern of negative deviations spanning parietal, occipital, and temporal regions, with relative positive deviations in right parietal and precentral regions. The partially overlapping but distinct patterns suggest that SP volume and thickness capture complementary aspects of development, potentially reflecting different processes underlying regional growth and cortical organization.^64^ Overall, the bidirectional and spatially structured deviations again argue against a single diffuse alteration in CHD.

Normative modelling therefore provided information not captured by group-level comparisons alone, particularly by identifying the extent and spatial organization of individual deviations. These estimates remain sensitive to normative cohort composition, segmentation uncertainty, and measurement noise and should be validated in larger cohorts.

### 4.4. Contribution of altered fetal hemodynamics and substrate delivery

Rising cerebral metabolic and substrate demand during the second and third trimesters^18,24^, together with experimental evidence for SP vulnerability to hypoxic-ischemic injury^34,65,66^ provide biological plausibility for a relationship between substrate delivery and SP development. We therefore examined whether substrate delivery scores proposed previously in CHD^8^, and CPR, a measure of relative cerebroplacental hemodynamics, might relate to SP deviations. Substrate delivery score was not independently associated with SP deviations. In contrast, higher CPR was associated with lower whole-brain SP thickness deviations, with the association strengthening after adjustment for overall brain size and inclusion of substrate delivery score, suggesting the two measures capture partially distinct aspects of the fetal circulatory environment. Lower CPR is generally interpreted as reflecting cerebrovascular redistribution (“brain sparing”) in response to physiological stress^27,67^. In our analyses, lower CPR was associated with less-negative SP thickness deviations, which could potentially reflect relative preservation of SP thickness in association with greater cerebrovascular redistribution, particularly after accounting for overall brain size. The specificity of these associations is, however, limited by the indirect nature and partial collinearity of these measures, as well as the multifactorial influences of genetic, placental, and cardiac physiology on fetal brain development in CHD.^68^ More direct measures of cerebral perfusion and oxygenation will be needed to determine whether these findings reflect a direct relationship between fetal cerebral hemodynamics and SP development.

### 4.5. Prenatal SP deviations and two-year neurodevelopmental outcomes

The SP is an active scaffold for subsequent brain development such that disturbances during this period may have lasting consequences.^31^ Children with CHD show prominent executive and language impairments^2^, and temporal and frontal SP volume deviations observed here overall align with the future substrates for executive and auditory-language networks^69,70^, as well as with preferential frontal and temporal SP alterations in CHD.^4^ Whole-brain SP volume deviations were associated with receptive language and fine motor scores, while right precuneus SP thickness remained associated with receptive and expressive communication after multiple-comparison correction. Although these associations were modest and warrant validation in larger longitudinal cohorts, they suggest that fetal SP deviations may have prognostic relevance.

### 4.6. Study considerations and future directions

Several considerations are important when interpreting these findings. The cross-sectional nature of the fetal MRI data limits inference about individual developmental trajectories and prevents determining whether observed SP deviations represent transient alterations or persistent developmental differences. The sample size, particularly for the hemodynamic and postnatal outcome analyses, also limits precision and statistical power for regional and interaction effects. Finally, the observational design and indirect characterization of fetal physiology preclude causal inference regarding the mechanisms linking CHD to SP development. These considerations motivate longitudinal imaging, larger cohorts, and more direct characterization of cerebral perfusion and oxygenation in future studies.

## 5. Conclusion

Fetuses with CHD show altered subplate development partially mirroring global reductions in brain growth, with superimposed spatially heterogeneous and individually variable regional deviations persisting beyond global scaling, particularly in posterior parietal, occipital, and temporal cortex. Normative modelling revealed bidirectional, spatially non-uniform deviations across individuals, consistent with heterogeneous developmental trajectories interacting with regionally distinct maturational programs. Associations with fetal hemodynamics and postnatal outcomes identify potential links between prenatal subplate development, fetal physiology, and later neurodevelopment, although these findings remain exploratory. Longitudinal imaging, refined circulatory phenotyping, and long-term neurodevelopmental follow-up will be needed to determine their developmental persistence and clinical significance.

## Data availability

Image processing code is available on GitHub (https://github.com/FNNDSC). Other processing used publicly available toolboxes detailed in *Materials and Methods*. Analyses were done mainly in Python (3.11.2) and RStudio (4.4.3). Manual corrections and some visualizations used Freeview (3.0). Anonymized MRI data may be shared with qualified researchers upon reasonable request and completion of a data transfer or data use agreement.

## Acknowledgements

We are deeply grateful to the families whose participation made this research possible. We thank Henry A. Feldman for his thoughtful feedback and insightful discussion regarding our analyses.

## Abbreviations

2V: two-ventricle
AIC: Akaike Information Criterion
ANCOVA: analysis of covariance
BIC: Bayesian Information Criterion
CHD: congenital heart disease
CP: cortical plate
CPR: cerebroplacental ratio
ECHO: echocardiogram
FAS: fetal aortic stenosis
FDR: false discovery rate
GA: gestational age
HASTE: half-Fourier single-shot turbo spin echo
HLHS: hypoplastic left heart syndrome
IZ: intermediary zone
MCA PI: middle cerebral artery pulsatility index
SP: subplate
SV: single-ventricle
TD: typically developing
TE: echo time
TGA: transposition of the great arteries
TOF: tetralogy of Fallot
TR: repetition time
UA PI: umbilical artery pulsatility index.

