## Supplementary Materials for "Fetal MRI Reveals Altered Subplate Development in Congenital Heart Disease"

### Supplementary Information (SI)

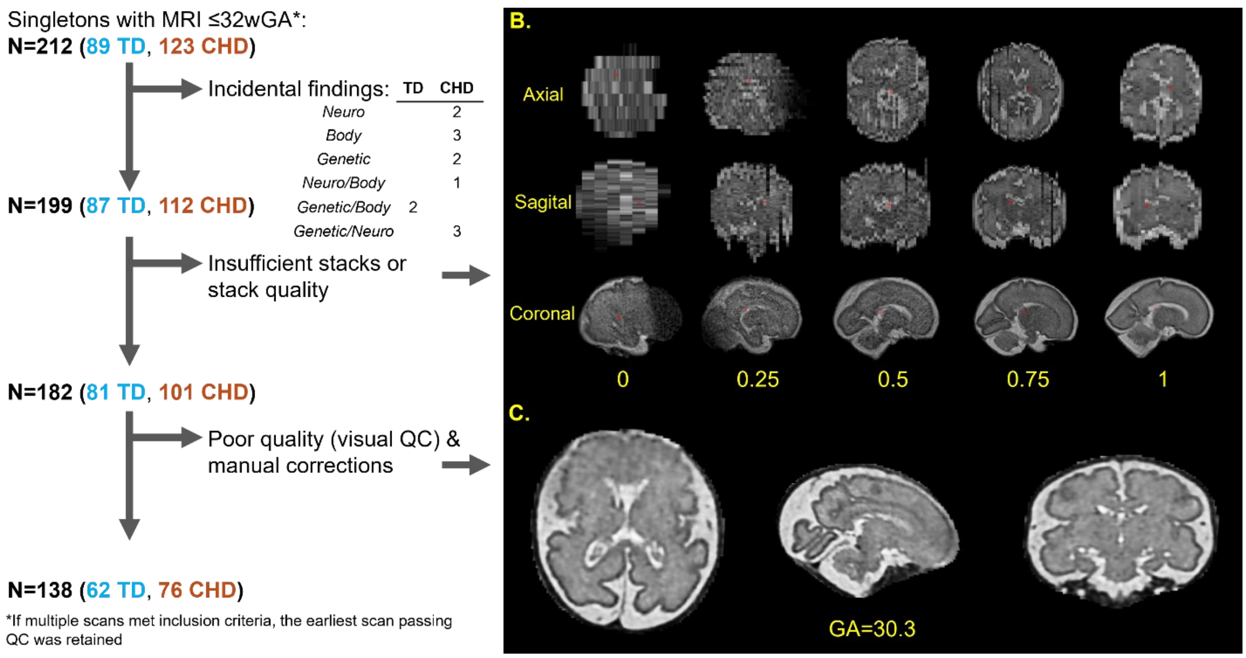


**SI-Fig1A.** Overview of the data inclusion and quality*. **B.** Illustration of the automated HASTE stack quality assessment. Before reconstruction, each available HASTE stack was assigned a value between 0-1 (0: very poor, unusable due to severe motion/blurring; 0.25: poor, usable only for basic reconstruction with evident artifacts; 0.5: acceptable, some blurring but preserved anatomical detail; 0.75: good, minimal blurring, clear tissue boundaries; 1: excellent, sharp anatomical detail, negligible motion artifacts) using a deep-learning model trained on slice-level manual quality ratings from two expert readers as ground truth. Only the subjects with at least 3 stacks with quality ≥0.4 were subsequently reconstructed with NeSVoR (this threshold achieves that stacks with excessive motion and generally low quality do not prevent effective reconstruction, NB: we have included one TD subjects with 2 stacks due to their very high quality). **C.** Example subject excluded after reconstruction due to insufficient reconstruction quality preventing robust delineation of the subplate.

*Overall inclusion rates did differ significantly between groups (TD: 62/89=70%; CHD: 76/123=62%; Fisher's exact test: OR=1.42, p=0.240), neither did the post-reconstruction QC-based exclusion (TD: 19/81=23%; CHD: 25/101=25%, Fisher's exact test: OR=0.93, p=0.760). The mean stack quality did not differ between TD and CHD groups (TD: 0.77 [0.54-0.91] mean [range]; CHD: 0.75 [0.52-0.92] mean [range]; Welch's t-test: t=-1.43, p=0.155, Cohen's D=0.24). Groups also showed comparable stack numbers (TD: 15 [2-32] mean [range]; CHD: 16 [4-25] mean [range]; Welch's t-test: t=1.09, p=0.277, Cohen's D=0.19).

Additionally, inclusion rates were similar across CHD subgroups (HLHS/TGA: 30/37=81%; Other-CHD: 46/64=72%, Fisher's exact test: OR=1.68, p=0.320), and more granular diagnosis (HLHS: 12/15=80%; TGA: 18/21=86%; TOF: 14/19=74%; FAS: 7/10=70%; other two ventricle: 16/23=70%; other single ventricle: 9/11=82%; likelihood ratio test, $\chi^2$=1.01, p=0.960), providing no evidence of preferential exclusion of any diagnostic category.

**A.**

**B.**

| | β | Z | p |
|---|---|---|---|
| **GA** | -0.001 | -3.35 | 0.001 |

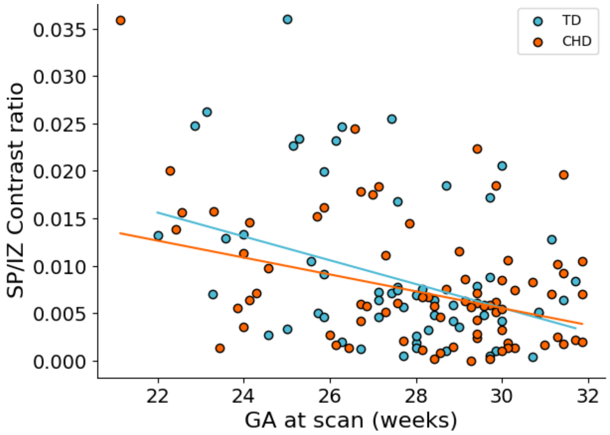


| | | | |
|---|---|---|---|
| **group** | 0.008 | 0.66 | 0.512 |
| **GA:group** | -0.000 | -0.62 | 0.533 |

**SI-Fig2A.** Scatterplot of SP/IZ contrast ratio as a function of GA at scan (colour coded by group belonging). The contrast ratio was calculated as: (mean SP signal − mean IZ signal) / (mean SP signal + mean IZ signal). The mean IZ signal was computed across voxels defined by expanding the SP tissue masks by 3 voxels into the IZ tissue. Solid lines represent robust linear regression fits for each group. **B.** Robust linear models (M-estimator) were used to examine the effects of GA, group (TD vs CHD) and their interaction SP/IZ contrast ratio. Estimates (β), z-statistics, and p-values are reported.

**SI-Table1A.** Linear regression models examining group differences (TD vs CHD) in neurodevelopmental outcomes, adjusting for Bayley edition and age at Bayley assessment. **B.** Linear regression models examining group differences in Doppler measures, adjusting for ECHO-MRI delay. In both cases, model effects are summarized using Type II analysis of variance (ANOVA). Reported are F-statistics (F), p-values (p), and partial eta squared (η²).

**A.**

| ***Neurodevelopmental outcomes:*** | **F** | **p** | **η²** |
|---|---|---|---|
| Gross motor: | | | |
| *group* | 10.80 | 0.002 | 0.12 |
| *Bayley edition* | 10.67 | 0.002 | 0.12 |
| *age at Bayley* | 0.35 | 0.557 | 0.00 |
| Fine motor: | | | |
| *Group* | 2.51 | 0.117 | 0.03 |
| *Bayley edition* | 1.26 | 0.265 | 0.02 |
| *age at Bayley* | 3.02 | 0.086 | 0.04 |
| Expressive language: | | | |
| *group* | 2.45 | 0.122 | 0.03 |
| *Bayley edition* | 3.47 | 0.066 | 0.04 |
| *age at Bayley* | 2.53 | 0.116 | 0.03 |
| Receptive language: | | | |
| *group* | 3.37 | 0.070 | 0.04 |
| *Bayley edition* | 2.58 | 0.113 | 0.03 |
| *age at Bayley* | 0.02 | 0.901 | 0.00 |
| Cognitive: | | | |
| *group* | 6.77 | 0.011 | 0.08 |
| *Bayley edition* | 8.40 | 0.005 | 0.10 |
| *age at Bayley* | 1.92 | 0.170 | 0.02 |

**B.**

| ***Doppler measures:*** | **F** | **p** | **η²** |
|---|---|---|---|
| MCA PI (Z): | | | |
| *group* | 1.17 | 0.287 | 0.04 |
| *ECHO-MRI delay* | 0.01 | 0.910 | 0.00 |
| *interaction* | 0.45 | 0.509 | 0.01 |
| UA PI (Z): | | | |
| *Group* | 0.53 | 0.468 | 0.01 |
| *ECHO-MRI delay* | 0.46 | 0.498 | 0.01 |
| *interaction* | 1.44 | 0.234 | 0.02 |
| CPR (Z): | | | |
| *group* | 1.22 | 0.296 | 0.11 |
| *ECHO-MRI delay* | 1.37 | 0.269 | 0.12 |
| *interaction* | 0.65 | 0.438 | 0.06 |

**SI-Table2.** Expanded subject characteristics for TD and CHD groups. Differences between groups for continuous variables were assessed using t-tests with Welch’s correction for unequal variances (t: t-statistic, dof: degrees of freedom, p: p-value, D: Cohen’s D). For categorical variables, differences were evaluated using Chi-squared (X²) tests (X²: X²-statistic, V: Cramer’s V). N: number of subjects.

| | TD (N=62) | CHD (N=76) | |
|---|---|---|---|
| ***Fetal characteristics:*** | | | |
| Fetal weight (percentile): median [range] (N) | 27.8 [0.0-93.9] (37) | 62.0 [0.0-100.0] (54) | t=-3.82, dof=82.43, p<0.000, D=0.80 |
| Growth restriction: count/available (%) | 0/17 (0) | 4/41 (10) | X²=0.59, dof=1.00, p=0.444, V=0.10 |
| Umbilical artery pulsatility index (Z): median [range] | 0.3 [-0.7-3.1] (56) | 0.6 [-1.3-3.3] (72) | t=-1.86, dof=124.75, p=0.069, D=0.32 |
| UA PI abnormality (Z>1.65): count (%) | 4 (7) | 9 (13) | X²=0.49, dof=1.00, p=0.484, V=0.06 |
| Middle cerebral artery pulsatility index (Z): median [range] | -0.2 [-1.4-0.7] (57) | -0.4 [-1.6-1.1] (71) | t=1.42, dof=125.04, p=0.159, D=0.24 |
| MCA PI abnormality (Z<-1.65): count (%) | 0 (0) | 0 (0) | - |
| ***Birth characteristics:*** | | | |
| GA at birth (weeks): median [range] (N) | 39.4 [34.6-41.9] (62) | 39.0 [31.7-40.6] (74) | t=3.25, dof=133.48, p=0.001, D=0.55 |
| Prematurity categories: count (%) | 62 | 74 | X²=1.50, dof=2.00, p=0.472, V=0.11 |
| *Term: ≥ 37* | 57 (92) | 64 (87) | |
| *Moderate/late preterm: [32, 37)* | 5 (8) | 9 (12) | |
| *Very preterm: ≤ 32* | 0 (0) | 1 (1) | |
| Birthweight (kg) median [range] (N) | 3.4 [2.3-4.6] (61) | 3.2 [1.0-4.9] (70) | t=3.41, dof=128.92, p=0.001, D=0.59 |
| Birthweight (Z) median [range] (N) | 0.7 [-1.0-3.3] (58) | 0.6 [-2.5-3.5] (62) | t=1.35, dof=117.41, p=0.179, D=0.25 |
| Birth head circumference (cm) median [range] (N) | 34.5 [30.5-38.0] (57) | 33.5 [27.0-37.0] (67) | t=3.87, dof=120.00, p<0.000, D=0.69 |
| Birth head circumference (Z) median [range] (N) | 0.7 [-2.0-3.0] (55) | 0.1 [-1.8-2.5] (60) | t=2.65, dof=107.51, p=0.009, D=0.50 |
| Apgar 1 median [range] (N) | 8 [2-9] (58) | 8 [1-9] (65) | t=2.49, dof=120.73, p=0.014, D=0.45 |
| Apgar 5 median [range] (N) | 9 [7-10] (58) | 8 [4-9] (65) | t=4.88, dof=98.10, p<0.000, D=0.85 |
| Mode of delivery: count (%) | 61 | 68 | X²=0.48, dof=2.00, p=0.786, V=0.06 |
| *Vaginal* | 46 (76) | 50 (73) | |
| *Elective C-section* | 10 (16) | 10 (15) | |
| *Emergent/non-elective C-section* | 5 (8) | 8 (12) | |
| CICU/NICU stay: count/available (%) | 3/56 (5) | 69/69 (100) | X²=109.53, dof=1.00, p<0.000, V=0.94 |
| Hospital stay (days): median [range] | 2 [0-26] (57) | 21 [2-143] (67) | t=-7.30, dof=68.56, p<0.000, D=1.22 |
| ***Maternal characteristics:*** | | | |
| Maternal age (years): median [range] (N) | 32.8 [21.8-39.9] (62) | 31.6 [21.8-41.1] (76) | t=1.19, dof=130.32, p=0.236, D=0.20 |
| Maternal BMI: median [range] (N) | 25.8 [18.7-44.3] (56) | 25.6 [17.7-42.8] (71) | t=0.54, dof=110.67, p=0.591, D=0.10 |
| BMI categories: count (%) | 56 | 71 | X²=2.65, dof=3.00, p=0.449, V=0.14 |
| *Underweight: < 18.5* | 0 (0) | 2 (3) | |
| *Healthy weight: (18.5, 25]* | 26 (47) | 28 (39) | |
| *Overweight: (25, 30]* | 17 (30) | 27 (38) | |
| *Obesity: ≥ 30* | 13 (23) | 14 (20) | |
| Education[#]: count (%) | 62 | 75 | X²=2.70, dof=3.00, p=0.440, V=0.14 |
| *Primary* | 6 (10) | 6 (8) | |
| *Secondary* | 10 (16) | 18 (24) | |
| *Tertiary* | 18 (29) | 26 (35) | |
| *Graduate* | 28 (45) | 25 (33) | |
| Income[&]: count (%) | 61 | 74 | X²=1.28, dof=2.00, p=0.528, V=0.10 |
| *Low* | 10 (16) | 8 (11) | |
| *Middle* | 32 (53) | 45 (61) | |
| *High* | 19 (31) | 21 (28) | |

| | | | |
|---|---|---|---|
| Language: count (%) | 55 | 57 | |
| *Monolingual (English)* | 42 (76) | 44 (77) | X²=2.18, dof=2.00, p=0.337, V=0.14 |
| *Monolingual (other)* | 0 (0) | 2 (4) | |
| *Multilingual* | 13 (24) | 11 (19) | |
| ***Cardiac characteristics:*** | | | |
| Substrate delivery: count (%) | | | |
| *normal* | - | 25 (33) | |
| *mixed* | - | 31 (41) | |
| *low* | - | 20 (26) | |
| Retrograde aortic flow: count (%) | - | 17 (22) | |
| Aortic arch hypoplasia: count (%) | - | 31 (41) | |
| ***Known postnatal events:*** | | | |
| Catheterization: count (%) | - | 37/63 (59) | |
| Cardiac surgery: count (%) | - | 59/63 (94) | |
| Seizures: count (%) | - | 4/63 (6) | |
| Abnormal postnatal neuro*: count (%) | - | 11/63 (17) | |
| Death: count (%) | - | 7/63 (11) | |

[#]Education categories: primary (less than college), secondary (partial college, specialized training, high school), tertiary (graduated standard college, university), graduate (graduate degree). [&]Income categories: low (≤50,000$), middle (50,000$-150,000$), high (>150,000$).* including microhemorrhages, white matter injury, and embolic/ischemic stroke.

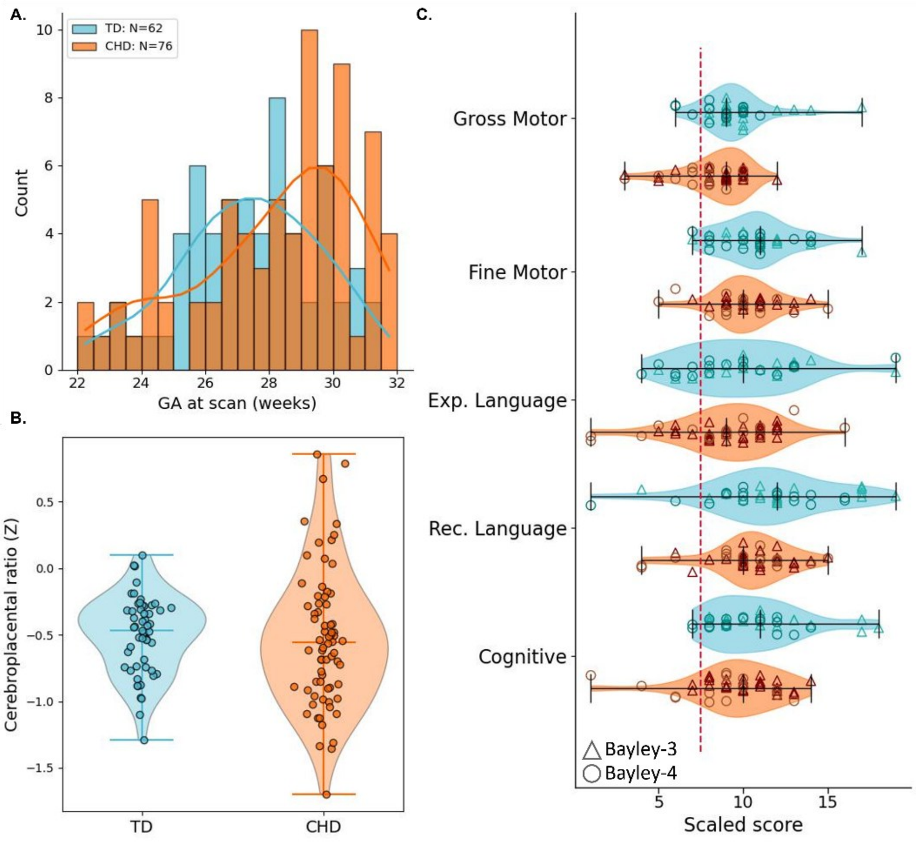


**SI-Fig3A.** Distribution of GA across TD and CHD groups. Overlaid lines indicate kernel density estimates scaled to histogram heights. **B.** Distribution of CPR (Z). **C.** Distribution of neurodevelopmental outcomes. TD group in blue, CHD group in orange.

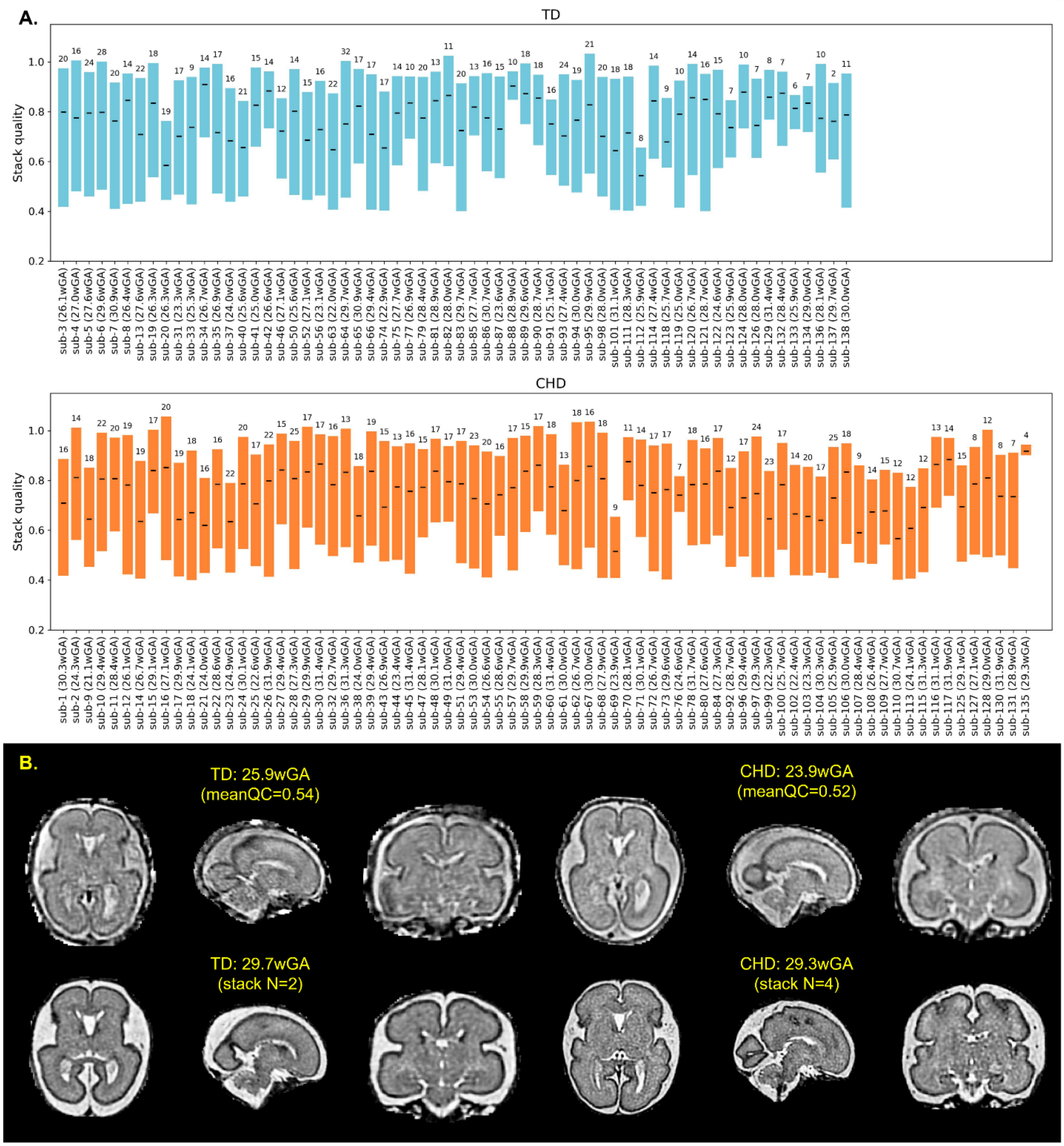


**SI-Fig4A.** Distribution of HASTE stack quality and numbers across included subjects, ordered by GA at scan. Black bars indicate each subject's mean stack quality, the boxes span the minimum–maximum quality range, and the number above each box shows the final number of stacks included in the subject's reconstruction. **B.** Example reconstructions for representative TD and CHD subjects. The upper row shows subjects with the lowest mean stack quality, and the lower row shows subjects with the fewest stacks included in the reconstruction.

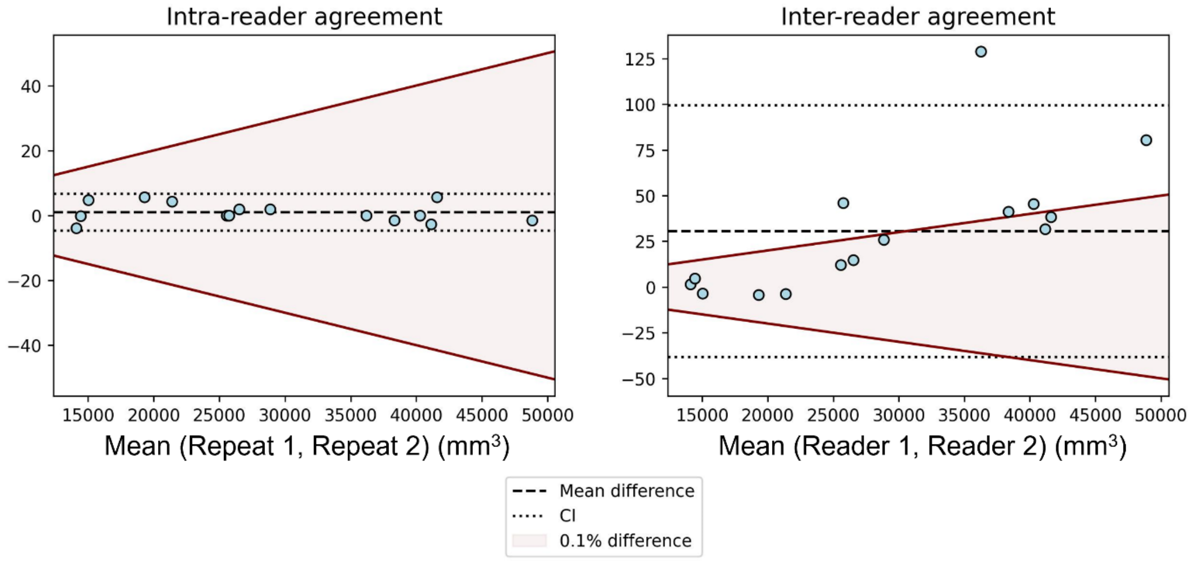


**SI-Fig5A.** Intra- and inter-reader reliability of SP segmentation visualized through Bland-Altman plots. The agreement was evaluated across 15 randomly selected subjects spanning the full range of gestational ages (mean GA 27.2 weeks, range 22.7–31.3). Intra-reader comparisons (left) show segmentations resulting from Reader 1's corrections at two repeats (Reader 1 corrected all data in this study). Inter-reader comparisons show segmentations from Reader 1 versus those from another trained reader (Reader 2). Results demonstrate high consistency of manual corrections. The red shaded area indicates a threshold of 0.1% difference relative to mean SP volume, showing that most inter-reader comparisons and all intra-reader comparisons fall within this range. High agreement was also observed when comparing segmentation overlaps (Dice coefficient): intra-reader mean ≈1, SD ≈0; inter-reader mean 0.99, SD ≈0.

**A.**

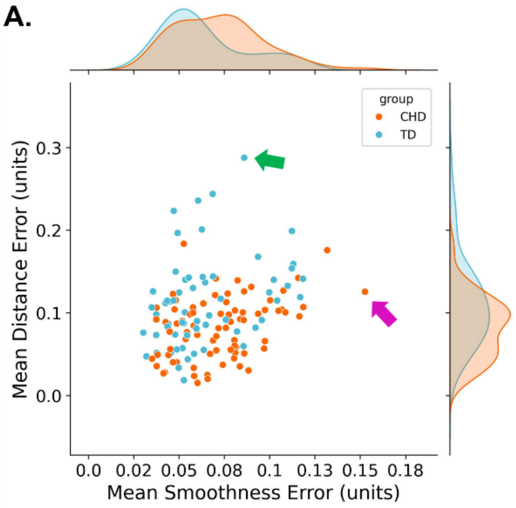


| | β | Z | p |
|---|---|---|---|
| **Smoothness error:** | | | |
| *GA* | 0.007 | 11.60 | <0.001 |
| *group* | -0.022 | -0.76 | 0.449 |
| *GA:group* | 0.001 | 0.65 | 0.519 |
| **Distance error:** | | | |
| *GA* | 0.003 | 1.89 | 0.059 |
| *group* | -0.077 | -0.98 | 0.326 |
| *GA:group* | 0.004 | 1.36 | 0.174 |

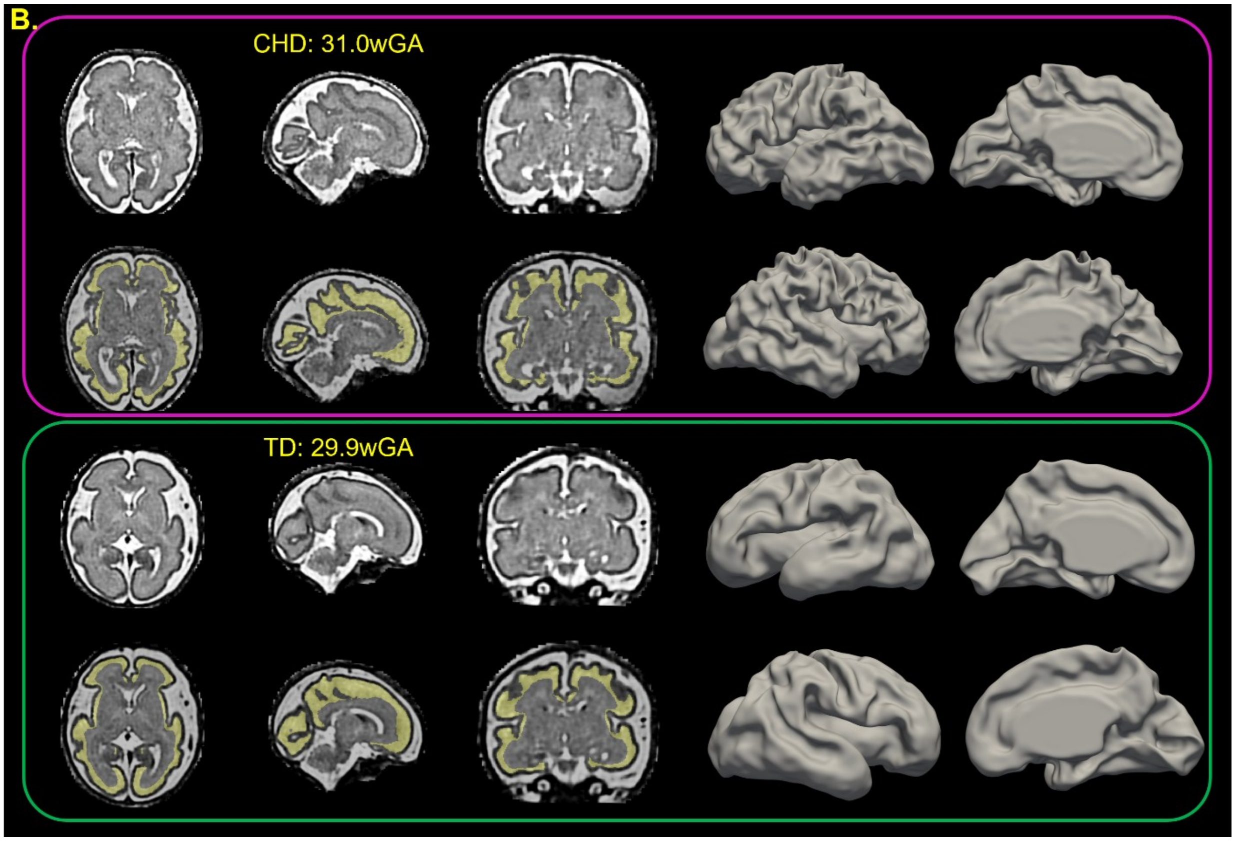


**SI-Fig6A.** Quality control of surface extraction for the outer SP surface across subjects. **B.** Robust linear models (M-estimator) were used to examine the effects of GA, group (TD vs CHD) and their interaction on surface errors. Estimates (β), z-statistics, and p-values are reported. **C.** Example T2w reconstruction with associated SP segmentation in yellow (lower left) in 31w template space for the subjects with the lowest surface extraction quality, as assessed by smoothness (in purple box) and distance errors (in green box). Corresponding outer SP surfaces for these subjects are shown on the right.

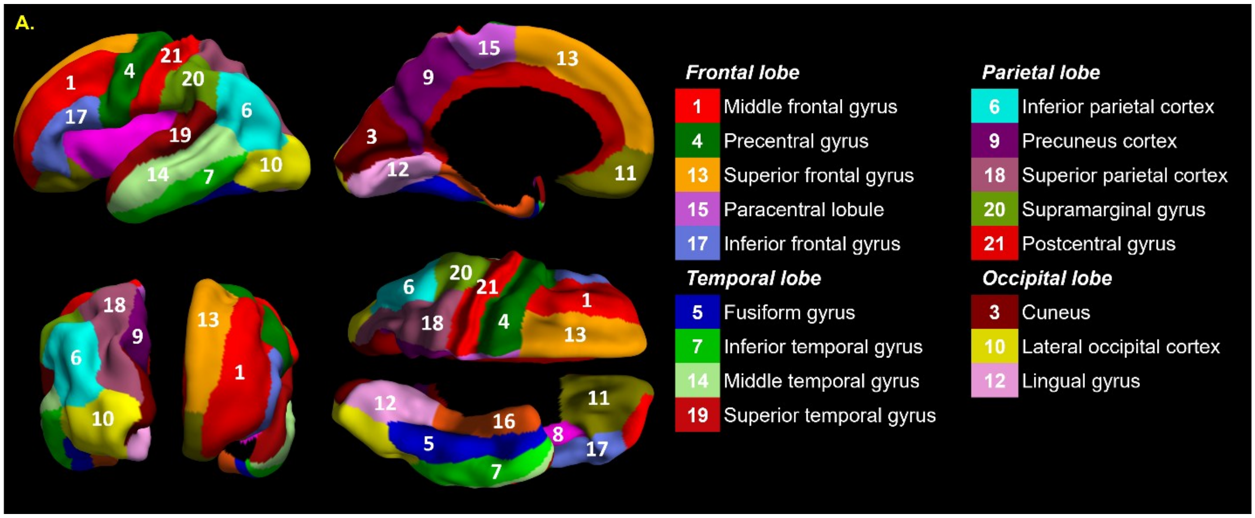

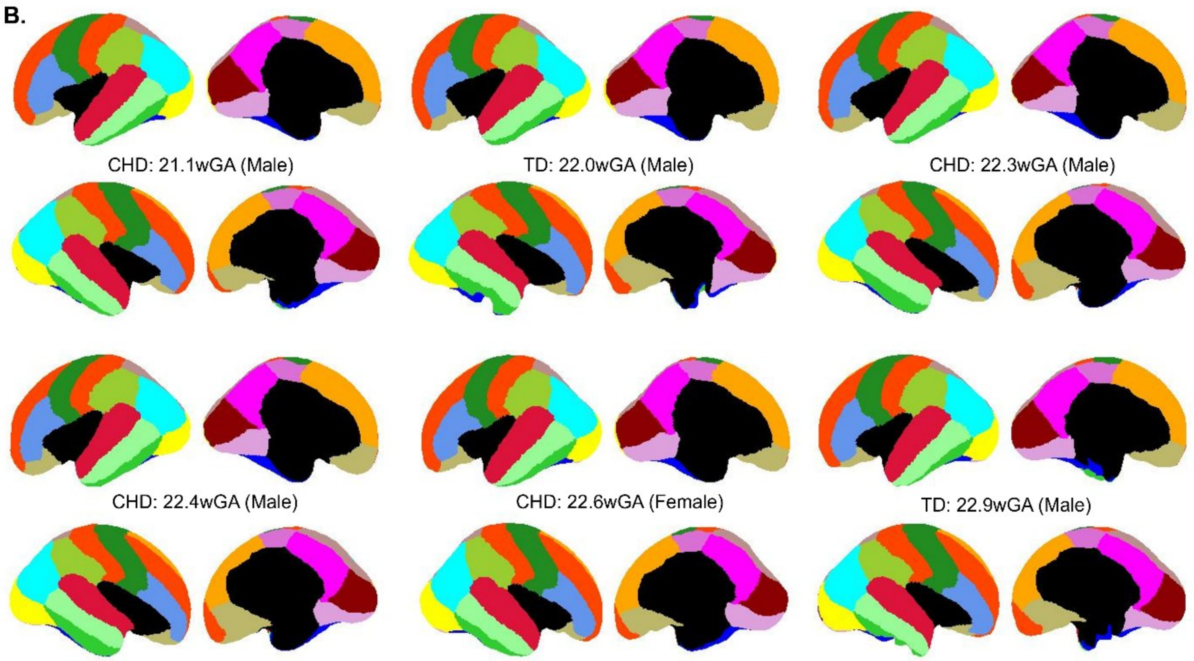


**SI-Fig7A.** Visualization of the final set of regions used in the study, along with the corresponding color scheme and lobar assignments. **B.** Example surface parcellations for the six youngest subjects from CHD and TD groups.

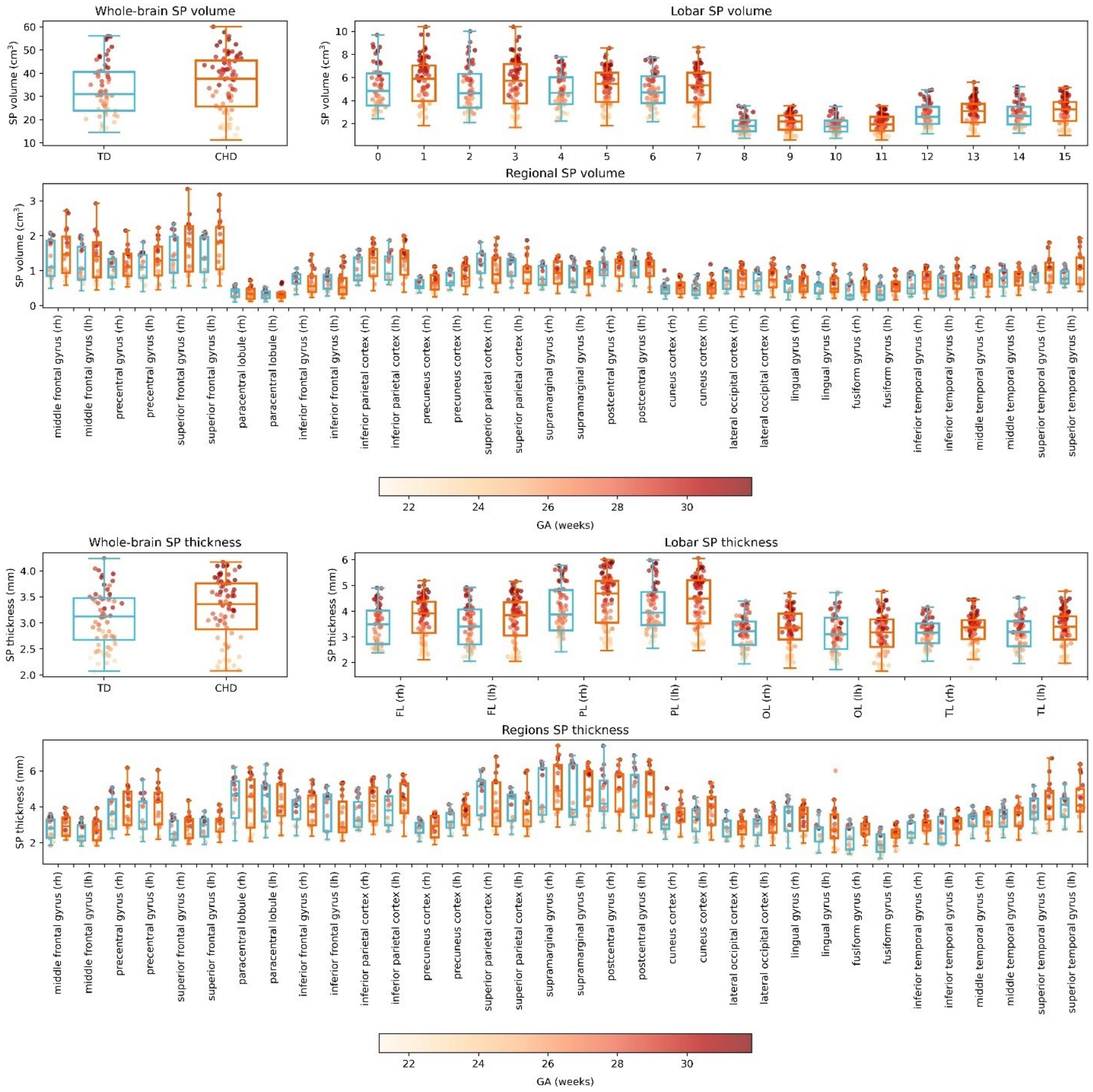


**SI-Fig8A.** Raw SP volume and thickness across whole-brain, lobar, and regional levels. Boxplots visualize the two groups (TD: blue, CHD:orange) at each scale; points show individual subjects colored by gestational age (GA, weeks), illustrating developmental variation.

**SI-Table3.** Linear mixed-effects models testing associations between SP thickness and SP depth at different spatial levels (A: hemisphere, B: lobe, C: region). SP depth was adjusted for sex and GA using a third-order spline prior to analysis and decomposed into between-subject and within-subject components. All models included GA modeled with a fourth-order spline. Effects were evaluated using Type II analysis of variance (ANOVA). Reported statistics include F-values, p-values (Pr(>F)), and partial eta-squared ($\eta^2$).

| **A.** | **SP thickness: hemisphere** | | |
|---|---|---|---|
| | **F** | **p** | **partial $\eta^2$** |
| **GA** | 1164.29 | <0.001 | 0.90 |
| **sex** | 8.58 | 0.003 | 0.06 |

| | | | |
|---|---|---|---|
| **Between-subject SP depth** | 0.68 | 0.411 | 0.00 |
| **Within-subject SP depth** | 25.82 | <0.001 | 0.00 |
| **hemi** | 6.94 | 0.008 | 0.00 |
| **Within-subject SP depth: hemi** | 0.93 | 0.335 | 0.00 |
| **B.** | **SP thickness: lobes** | | |
| | **F** | **p** | **partial η²** |
| **GA** | 1138.31 | <0.001 | 0.90 |
| **sex** | 8.93 | 0.002 | 0.06 |
| **Between-subject SP depth** | 0.22 | 0.638 | 0.00 |
| **Within-subject SP depth** | 23.14 | <0.001 | 0.00 |
| **hemi** | 139.55 | <0.001 | 0.03 |
| **lobe** | 1704.94 | <0.001 | 0.30 |
| **Within-subject SP depth: lobe** | 494.25 | <0.001 | 0.10 |
| **C.** | **SP thickness:region** | | |
| | **F** | **p** | **partial η²** |
| **GA** | 1124.01 | <0.001 | 0.90 |
| **sex** | 8.31 | 0.004 | 0.06 |
| **Between-subject SP depth** | 0.29 | 0.589 | 0.00 |
| **Within-subject SP depth** | 34.13 | <0.001 | 0.01 |
| **hemi** | 1003.54 | <0.001 | 0.14 |
| **region** | 18195.25 | <0.001 | 0.65 |
| **Within-subject SP depth: region** | 379.21 | <0.001 | 0.08 |

**A.**

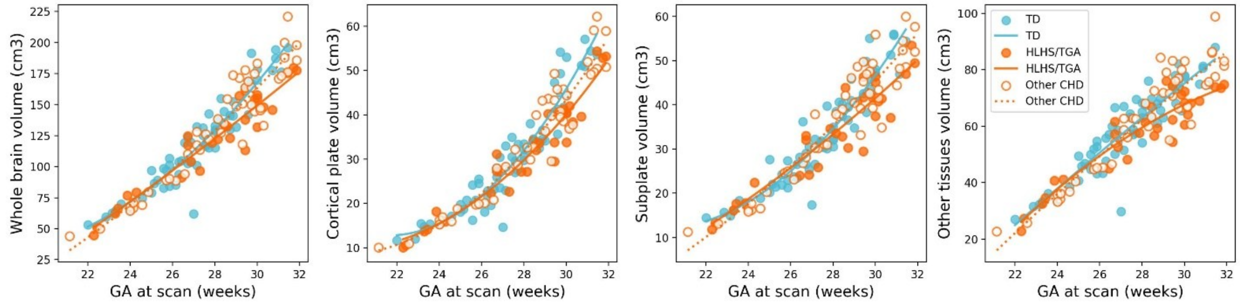


**B.**

| | **Whole brain volume** | | | **CP volume** | | | **SP volume** | | | **Other volume** | | |
|---|---|---|---|---|---|---|---|---|---|---|---|---|
| | **F** | **p** | **η²** | **F** | **p** | **η²** | **F** | **p** | **η²** | **F** | **p** | **η²** |
| **subgroup** | 7.40 | <0.001 | 0.10 | 5.83 | 0.004 | 0.08 | 4.51 | 0.013 | 0.06 | 7.73 | <0.001 | 0.10 |
| **sex** | 21.06 | <0.001 | 0.10 | 8.51 | 0.004 | 0.04 | 19.76 | <0.001 | 0.11 | 22.10 | <0.001 | 0.11 |
| **GA** | 833.78 | <0.001 | 0.93 | 1692.71 | <0.001 | 0.93 | 736.23 | <0.001 | 0.92 | 567.08 | <0.001 | 0.90 |

| | **Whole brain volume** | | **CP volume** | | **SP volume** | | **Other volume** | |
|---|---|---|---|---|---|---|---|---|
| | **t** | **q** | **t** | **q** | **t** | **q** | **t** | **q** |

| | | | | | | | | |
|---|---|---|---|---|---|---|---|---|
| TD vs HLHS/TGA | 3.79 | <0.001 | 3.41 | 0.003 | 2.89 | 0.014 | 3.87 | <0.001 |
| TD vs Other CHD | 0.90 | 0.368 | 1.05 | 0.294 | 0.40 | 0.692 | 0.92 | 0.361 |
| HLHS/TGA vs Other CHD | -2.92 | 0.006 | -2.39 | 0.027 | -2.47 | 0.022 | -2.99 | 0.005 |

**SI-Fig9A.** Unadjusted whole brain and tissue volumes stratified by group: TD (blue), CHD subgroups HLHS/TGA (orange), and Other CHD (white with orange border). Lines represent the best-fitting model (linear or quadratic), selected using the Akaike Information Criterion (AIC) and estimated using a RANSAC algorithm to visualize relationships with gestational age (GA). **B.** Results from linear regression (ANCOVA) models assessing group differences in log-transformed whole brain and tissue volumes, adjusting for GA, and sex. Model effects are summarized using Type II analysis of variance (ANOVA). Reported are F-statistics (F), p (PR(>F)), and partial eta squared ($\eta^2$). Post hoc pairwise comparisons between groups were performed using covariate-adjusted linear contrasts derived from the fitted ANCOVA models (q: FDR-corrected p-values).

**SI-Table4A.** Type II analysis of variance for the linear model predicting log-transformed SP volume, adjusting for GA (modeled using a cubic spline with 2 degrees of freedom) and sex. Reported statistics include F-statistics, p-values (p), and partial eta-squared ($\eta^2$). **B.** Type III analysis of variance for mixed-effects models predicting log-transformed SP volume, adjusting for GA (cubic spline, 2 degrees of freedom) and sex. For each tissue type, lobe, or region, adjusted group means for CHD and TD are presented (evaluated at mean GA = 27.8 weeks, mode sex = male). Effect sizes are reported as Cohen's d, and p-values correspond to FDR-corrected significance tests (q, across lobes for lobar analyses and across regions for regional analyses). Significant group differences are color-coded for visualization, with blue indicating TD > CHD and orange indicating CHD > TD. **C.** As in A, with models additionally adjusting for residual brain volume (adjusted for linear effects of GA and sex). **D.** As in B, with models additionally adjusting for residual whole-brain SP volume (adjusted for sex and GA using a second-degree spline). Group contrasts are evaluated at mean GA = 27.8 weeks, male sex, and mean SP volume = 34.26 cm³.

| **A.** | **SP volume** | | |
|---|---|---|---|
| | **F** | **p** | **partial $\eta^2$** |
| **GA** | 713.32 | <0.001 | 0.91 |
| **sex** | 15.98 | <0.001 | 0.12 |
| **group** | 2.82 | 0.096 | 0.02 |

| **B.** | ***Left SP volume (cm³)*** | | | | ***Right SP volume (cm³)*** | | | |
|---|---|---|---|---|---|---|---|---|
| | **CHD** | **TD** | **D** | **q** | **CHD** | **TD** | **D** | **q** |
| **Global** | 14.19 | 14.88 | -1.69 | 0.024 | 14.4 | 14.89 | -1.19 | 0.111 |
| ***Frontal*** | 4.90 | 5.06 | -0.57 | 0.139 | 5.06 | 5.17 | -0.36 | 0.348 |
| Middle frontal gyrus | 1.26 | 1.29 | -0.20 | 0.415 | 1.4 | 1.39 | 0.04 | 0.883 |
| Precentral gyrus | 1.16 | 1.27 | -0.71 | 0.003 | 1.12 | 1.21 | -0.67 | 0.006 |
| Superior frontal gyrus | 1.51 | 1.53 | -0.13 | 0.589 | 1.51 | 1.51 | -0.01 | 0.982 |
| Paracentral lobule | 0.32 | 0.33 | -0.43 | 0.078 | 0.35 | 0.36 | -0.39 | 0.107 |
| Inferior frontal gyrus | 0.64 | 0.63 | 0.06 | 0.816 | 0.67 | 0.68 | -0.09 | 0.700 |
| ***Parietal*** | 4.72 | 5.08 | -1.23 | 0.001 | 4.75 | 5.00 | -0.87 | 0.023 |
| Inferior parietal cortex | 1.15 | 1.25 | -0.74 | 0.002 | 1.15 | 1.19 | -0.28 | 0.251 |

| | | | | | | | | |
|---|---|---|---|---|---|---|---|---|
| Precuneus cortex | 0.78 | 0.85 | -0.82 | 0.001 | 0.64 | 0.71 | -0.97 | <0.001 |
| Superior parietal cortex | 0.94 | 0.99 | -0.48 | 0.049 | 1.03 | 1.06 | -0.22 | 0.358 |
| Supramarginal gyrus | 0.82 | 0.87 | -0.48 | 0.048 | 0.85 | 0.89 | -0.38 | 0.124 |
| Postcentral gyrus | 1.01 | 1.07 | -0.53 | 0.030 | 1.06 | 1.12 | -0.52 | 0.033 |
| ***Occipital*** | 1.78 | 1.88 | -0.92 | 0.017 | 1.89 | 1.9 | -0.10 | 0.794 |
| Cuneus | 0.50 | 0.56 | -0.87 | <0.001 | 0.52 | 0.52 | 0.00 | 0.989 |
| Lateral occipital cortex | 0.75 | 0.74 | 0.12 | 0.625 | 0.77 | 0.77 | 0.04 | 0.879 |
| Lingual gyrus | 0.52 | 0.57 | -0.87 | <0.001 | 0.59 | 0.6 | -0.20 | 0.421 |
| ***Temporal*** | 2.78 | 2.86 | -0.46 | 0.229 | 2.68 | 2.81 | -0.79 | 0.039 |
| Fusiform gyrus | 0.44 | 0.46 | -0.42 | 0.084 | 0.47 | 0.48 | -0.10 | 0.679 |
| Inferior temporal gyrus | 0.64 | 0.65 | -0.12 | 0.632 | 0.64 | 0.65 | -0.13 | 0.599 |
| Middle temporal gyrus | 0.78 | 0.81 | -0.40 | 0.102 | 0.7 | 0.75 | -0.59 | 0.016 |
| Superior temporal gyrus | 0.90 | 0.91 | -0.12 | 0.613 | 0.86 | 0.92 | -0.64 | 0.009 |

| **C.** | **SP volume** | | |
|---|---|---|---|
| | **F** | **p** | **partial $\eta^2$** |
| **GA** | 1898.93 | <0.001 | 0.98 |
| **sex** | 34.12 | <0.001 | 0.21 |
| **residual brain volume** | 408.02 | <0.001 | 0.76 |
| **group** | 0.48 | 0.492 | 0.00 |

| **D.** | ***Left SP volume*** | | | | ***Right SP volume*** | | | |
|---|---|---|---|---|---|---|---|---|
| | **CHD** | **TD** | **D** | **q** | **CHD** | **TD** | **D** | **q** |
| **Global** | 14.47 | 14.66 | -0.52 | 0.004 | 14.68 | 14.67 | 0.03 | 0.882 |
| ***Frontal*** | 4.99 | 4.99 | 0.01 | 0.967 | 5.16 | 5.10 | 0.22 | 0.209 |
| Middle frontal gyrus | 1.28 | 1.27 | 0.09 | 0.580 | 1.43 | 1.37 | 0.33 | 0.054 |
| Precentral gyrus | 1.19 | 1.25 | -0.42 | 0.014 | 1.14 | 1.19 | -0.38 | 0.025 |
| Superior frontal gyrus | 1.54 | 1.51 | 0.16 | 0.345 | 1.54 | 1.49 | 0.29 | 0.092 |
| Paracentral lobule | 0.32 | 0.33 | -0.14 | 0.422 | 0.35 | 0.36 | -0.10 | 0.556 |
| Inferior frontal gyrus | 0.65 | 0.62 | 0.35 | 0.040 | 0.68 | 0.67 | 0.20 | 0.243 |
| ***Parietal*** | 4.81 | 5.00 | -0.67 | <0.001 | 4.84 | 4.93 | -0.30 | 0.082 |
| Inferior parietal cortex | 1.17 | 1.23 | -0.45 | 0.008 | 1.18 | 1.17 | 0.01 | 0.940 |
| Precuneus cortex | 0.79 | 0.84 | -0.53 | 0.002 | 0.65 | 0.70 | -0.68 | <0.001 |
| Superior parietal cortex | 0.96 | 0.98 | -0.19 | 0.274 | 1.05 | 1.04 | 0.07 | 0.686 |
| Supramarginal gyrus | 0.84 | 0.86 | -0.19 | 0.268 | 0.87 | 0.87 | -0.08 | 0.628 |
| Postcentral gyrus | 1.03 | 1.06 | -0.24 | 0.165 | 1.08 | 1.11 | -0.23 | 0.183 |
| ***Occipital*** | 1.81 | 1.85 | -0.35 | 0.043 | 1.92 | 1.87 | 0.48 | 0.006 |
| Cuneus | 0.51 | 0.55 | -0.58 | 0.001 | 0.53 | 0.51 | 0.30 | 0.082 |
| Lateral occipital cortex | 0.77 | 0.73 | 0.41 | 0.016 | 0.79 | 0.76 | 0.33 | 0.053 |
| Lingual gyrus | 0.53 | 0.56 | -0.58 | 0.001 | 0.6 | 0.59 | 0.1 | 0.571 |

| | | | | | | | | |
|---|---|---|---|---|---|---|---|---|
| ***Temporal*** | 2.83 | 2.81 | 0.11 | 0.509 | 2.74 | 2.77 | -0.22 | 0.196 |
| Fusiform gyrus | 0.45 | 0.45 | -0.13 | 0.453 | 0.48 | 0.47 | 0.19 | 0.260 |
| Inferior temporal gyrus | 0.65 | 0.64 | 0.18 | 0.302 | 0.65 | 0.64 | 0.17 | 0.334 |
| Middle temporal gyrus | 0.79 | 0.80 | -0.11 | 0.536 | 0.71 | 0.74 | -0.29 | 0.086 |
| Superior temporal gyrus | 0.91 | 0.90 | 0.17 | 0.320 | 0.87 | 0.91 | -0.35 | 0.041 |

**SI-Table5A.** Type II analysis of variance for the linear model predicting log-transformed SP thickness, adjusting for GA (modeled using a cubic spline with 4 degrees of freedom) and sex. Reported statistics include F-statistics, p-values, and partial eta-squared ($\eta^2$). **B.** Type III analysis of variance for mixed-effects models predicting log-transformed SP volume, adjusting for GA (cubic spline, 3 degrees of freedom), sex, and within-subject SP depth (based on results presented in SI-Table3). For each tissue type, lobe, or region, adjusted group means for CHD and TD are presented (evaluated at mean GA = 27.8 weeks, mode sex = male, and SP depth=6.08 mm). Effect sizes are reported as Cohen's d, and p-values correspond to FDR-corrected significance tests (q, across lobes for lobar analyses and across regions for regional analyses). Significant group differences are color-coded for visualization, with blue indicating TD > CHD and orange indicating CHD > TD. **C.** As in A, with models additionally adjusting for residual brain volume (adjusted for linear effects of GA and sex). **D.** As in B, with models additionally adjusting for residual whole-brain SP thickness (adjusted for sex and GA using a fourth-degree spline). Group contrasts are evaluated at mean GA = 27.8 weeks, male sex, and SP thickness = 3.33 mm, and SP depth=6.08 mm.

| **A.** | **SP thickness** | | |
|---|---|---|---|
| | **F** | **p** | **partial $\eta^2$** |
| **GA** | 212.64 | <0.001 | 0.87 |
| **sex** | 11.89 | <0.001 | 0.08 |
| **group** | 0.23 | 0.631 | 0.00 |

| **B.** | ***Left SP thickness (mm)*** | | | | ***Right SP thickness (mm)*** | | | |
|---|---|---|---|---|---|---|---|---|
| | **CHD** | **TD** | **D** | **q** | **CHD** | **TD** | **D** | **q** |
| **Global** | 3.51 | 3.62 | -0.69 | 0.041 | 3.61 | 3.62 | -0.10 | 0.777 |
| ***Frontal*** | 3.49 | 3.5 | -0.06 | 0.801 | 3.55 | 3.51 | 0.14 | 0.571 |
| Middle frontal gyrus | 2.49 | 2.57 | -0.40 | 0.146 | 2.75 | 2.72 | 0.18 | 0.516 |
| Precentral gyrus | 3.92 | 3.97 | -0.15 | 0.593 | 4.23 | 4.13 | 0.32 | 0.245 |
| Superior frontal gyrus | 2.81 | 2.86 | -0.26 | 0.349 | 2.68 | 2.68 | 0.02 | 0.936 |
| Paracentral lobule | 4.38 | 4.54 | -0.47 | 0.095 | 4.10 | 4.11 | -0.03 | 0.914 |
| Inferior frontal gyrus | 3.67 | 3.53 | 0.50 | 0.071 | 3.75 | 3.75 | 0.01 | 0.970 |
| ***Parietal*** | 4.22 | 4.29 | -0.23 | 0.356 | 4.32 | 4.24 | 0.25 | 0.296 |
| Inferior parietal cortex | 4.10 | 4.31 | -0.65 | 0.019 | 4.21 | 4.11 | 0.32 | 0.253 |
| Precuneus cortex | 3.66 | 3.93 | -0.91 | 0.001 | 3.34 | 3.55 | -0.80 | 0.004 |
| Superior parietal cortex | 3.81 | 3.89 | -0.27 | 0.333 | 4.23 | 4.04 | 0.58 | 0.035 |
| Supramarginal gyrus | 5.21 | 5.24 | -0.07 | 0.806 | 5.46 | 5.34 | 0.28 | 0.308 |
| Postcentral gyrus | 4.79 | 4.73 | 0.15 | 0.586 | 5.02 | 4.92 | 0.27 | 0.332 |
| ***Occipital*** | 2.91 | 3.12 | -0.97 | <0.001 | 3.12 | 3.15 | -0.16 | 0.517 |
| Cuneus | 3.31 | 3.62 | -1.17 | <0.001 | 3.25 | 3.33 | -0.32 | 0.251 |
| Lateral occipital cortex | 2.81 | 2.84 | -0.11 | 0.686 | 2.85 | 2.82 | 0.14 | 0.601 |

| | | | | | | | | |
|---|---|---|---|---|---|---|---|---|
| Lingual gyrus | 2.54 | 2.74 | -0.99 | <0.001 | 2.96 | 2.94 | 0.11 | 0.698 |
| ***Temporal*** | 3.22 | 3.33 | -0.42 | 0.089 | 3.26 | 3.35 | -0.38 | 0.122 |
| Fusiform gyrus | 2.27 | 2.44 | -0.93 | 0.001 | 2.59 | 2.72 | -0.63 | 0.023 |
| Inferior temporal gyrus | 2.92 | 2.96 | -0.18 | 0.526 | 2.82 | 2.89 | -0.31 | 0.264 |
| Middle temporal gyrus | 3.25 | 3.38 | -0.48 | 0.084 | 3.29 | 3.39 | -0.37 | 0.186 |
| Superior temporal gyrus | 4.28 | 4.39 | -0.30 | 0.273 | 4.43 | 4.50 | -0.21 | 0.458 |

| **C.** | **SP thickness** | | |
|---|---|---|---|
| | **F** | **p** | **partial $\eta^2$** |
| **GA** | 251.98 | <0.001 | 0.89 |
| **sex** | 11.16 | 0.001 | 0.08 |
| **residual brain volume** | 29.76 | <0.001 | 0.19 |
| **group** | 0.37 | 0.544 | 0.00 |

| **D.** | ***Left SP thickness (mm)*** | | | | ***Right SP thickness (mm)*** | | | |
|---|---|---|---|---|---|---|---|---|
| | **CHD** | **TD** | **D** | **q** | **CHD** | **TD** | **D** | **q** |
| **Global** | 3.53 | 3.60 | -0.62 | 0.001 | 3.62 | 3.61 | 0.07 | 0.710 |
| ***Frontal*** | 3.5 | 3.49 | 0.02 | 0.894 | 3.56 | 3.50 | 0.22 | 0.201 |
| Middle frontal gyrus | 2.47 | 2.53 | -0.24 | 0.166 | 2.71 | 2.66 | 0.20 | 0.255 |
| Precentral gyrus | 3.94 | 3.96 | -0.06 | 0.736 | 4.25 | 4.12 | 0.31 | 0.076 |
| Superior frontal gyrus | 2.79 | 2.84 | -0.16 | 0.355 | 2.67 | 2.65 | 0.06 | 0.710 |
| Paracentral lobule | 4.39 | 4.54 | -0.35 | 0.046 | 4.08 | 4.07 | 0.02 | 0.898 |
| Inferior frontal gyrus | 3.69 | 3.53 | 0.45 | 0.010 | 3.79 | 3.77 | 0.04 | 0.812 |
| ***Parietal*** | 4.25 | 4.29 | -0.15 | 0.39 | 4.36 | 4.25 | 0.34 | 0.053 |
| Inferior parietal cortex | 4.14 | 4.33 | -0.44 | 0.011 | 4.27 | 4.15 | 0.29 | 0.089 |
| Precuneus cortex | 3.65 | 3.89 | -0.64 | <0.001 | 3.31 | 3.51 | -0.57 | 0.001 |
| Superior parietal cortex | 3.85 | 3.93 | -0.19 | 0.280 | 4.22 | 4.01 | 0.51 | 0.003 |
| Supramarginal gyrus | 5.28 | 5.28 | 0.00 | 0.985 | 5.56 | 5.42 | 0.26 | 0.129 |
| Postcentral gyrus | 4.81 | 4.73 | 0.17 | 0.336 | 5.04 | 4.91 | 0.26 | 0.132 |
| ***Occipital*** | 2.93 | 3.13 | -0.92 | <0.001 | 3.14 | 3.17 | -0.10 | 0.585 |
| Cuneus | 3.33 | 3.64 | -0.88 | <0.001 | 3.28 | 3.35 | -0.19 | 0.268 |
| Lateral occipital cortex | 2.76 | 2.78 | -0.07 | 0.691 | 2.83 | 2.78 | 0.17 | 0.325 |
| Lingual gyrus | 2.55 | 2.74 | -0.71 | <0.001 | 2.97 | 2.93 | 0.14 | 0.435 |
| ***Temporal*** | 3.24 | 3.33 | -0.36 | 0.045 | 3.28 | 3.35 | -0.3 | 0.082 |
| Fusiform gyrus | 2.26 | 2.41 | -0.65 | <0.001 | 2.58 | 2.69 | -0.42 | 0.016 |
| Inferior temporal gyrus | 2.99 | 3.00 | -0.01 | 0.953 | 2.92 | 2.97 | -0.17 | 0.339 |
| Middle temporal gyrus | 3.22 | 3.32 | -0.3 | 0.078 | 3.31 | 3.38 | -0.21 | 0.230 |
| Superior temporal gyrus | 4.30 | 4.37 | -0.18 | 0.303 | 4.44 | 4.48 | -0.10 | 0.553 |

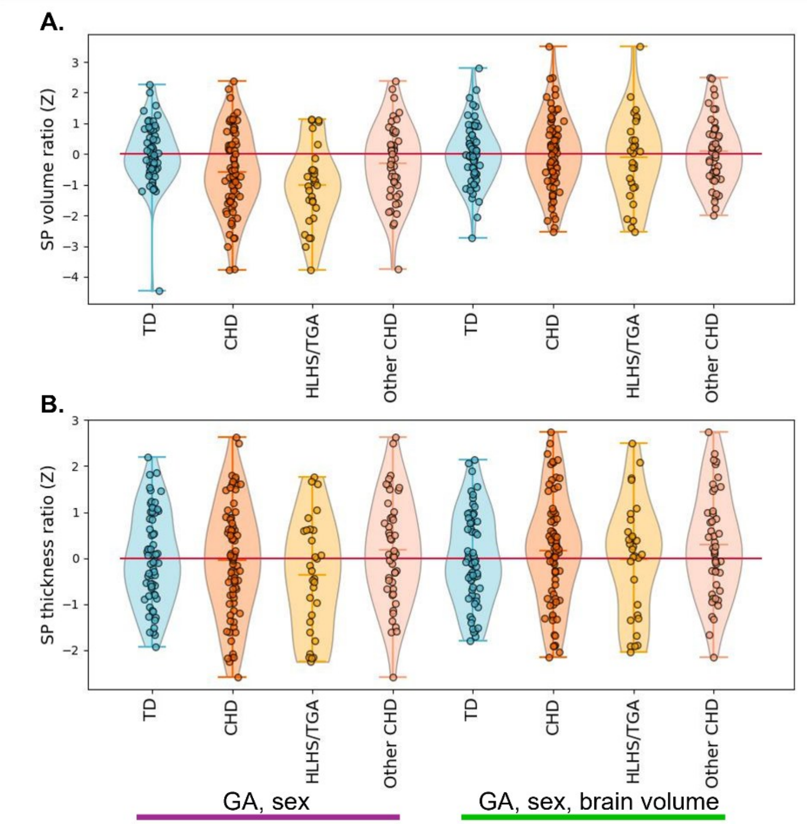


**SI-Fig10A.** Whole-brain SP volume deviations (Z-scores) for CHD subjects (and HLHS/TGA, Other-CHD subgroups) compared to TD reference. Points show individual subjects; red line indicates perfect agreement with the reference model. **B.** As in A but visualising SP thickness deviations. Colored-lines below figures indicate covariates used in the reference TD models (purple: GA and sex; green: GA, sex, and residualized brain volume).

**SI-Table6.** Post-hoc comparisons of CHD-associated deviations in SP volume (**A**) and SP thickness (**B**). Z-scores were compared to 0 using t-tests. Associated Cohen's D and FDR-corrected p-values (q) are reported.

| **A.** | ***Left SP volume*** | | | | ***Right SP volume*** | | | |
|---|---|---|---|---|---|---|---|---|
| | **Z ± SD** | **T** | **D** | **q** | **Z** | **T** | **D** | **q** |
| **Global** | -0.01±1.105 | -0.27 | 0.01 | 0.789 | 0.07±1.066 | 2.79 | 0.06 | 0.005 |
| ***Frontal*** | 0.09±1.149 | 1.81 | 0.07 | 0.071 | 0.12±1.101 | 2.64 | 0.10 | 0.008 |
| Middle frontal gyrus | 0.02±0.103 | 2.09 | 0.24 | 0.090 | 0.05±0.086 | 5.30 | 0.61 | <0.001 |
| Precentral gyrus | -0.04±0.099 | -3.12 | 0.36 | 0.008 | - 0.03±0.132 | -2.06 | 0.24 | 0.092 |
| Superior frontal gyrus | 0.03±0.113 | 2.51 | 0.29 | 0.035 | 0.05±0.133 | 3.10 | 0.36 | 0.008 |
| Paracentral lobule | -0.0±0.16 | -0.13 | 0.02 | 0.952 | 0.0±0.127 | 0.13 | 0.01 | 0.952 |
| Inferior frontal gyrus | 0.05±0.135 | 3.52 | 0.40 | 0.003 | 0.04±0.123 | 2.61 | 0.30 | 0.028 |
| ***Parietal*** | -0.13±1.008 | -2.72 | 0.10 | 0.007 | - 0.02±0.959 | -0.95 | 0.04 | 0.343 |
| Inferior parietal cortex | -0.04±0.099 | -3.43 | 0.39 | 0.003 | 0.02±0.081 | 1.62 | 0.19 | 0.154 |

| | | | | | | | | |
|---|---|---|---|---|---|---|---|---|
| Precuneus cortex | -0.05±0.096 | -4.33 | 0.50 | <0.001 | -0.07±0.108 | -5.29 | 0.61 | <0.001 |
| Superior parietal cortex | -0.01±0.123 | -0.58 | 0.07 | 0.659 | 0.02±0.105 | 1.81 | 0.21 | 0.126 |
| Supramarginal gyrus | -0.01±0.12 | -0.62 | 0.07 | 0.656 | 0.0±0.099 | 0.35 | 0.04 | 0.826 |
| Postcentral gyrus | -0.01±0.142 | -0.86 | 0.10 | 0.492 | -0.01±0.123 | -0.92 | 0.11 | 0.471 |
| ***Occipital*** | -0.08±1.173 | -1.38 | 0.07 | 0.169 | 0.20±1.047 | 3.96 | 0.19 | <0.001 |
| Cuneus | -0.05±0.134 | -3.52 | 0.40 | 0.003 | 0.05±0.123 | 3.43 | 0.39 | 0.003 |
| Lateral occipital cortex | 0.06±0.114 | 4.74 | 0.54 | <0.001 | 0.05±0.101 | 4.52 | 0.52 | <0.001 |
| Lingual gyrus | -0.05±0.133 | -3.54 | 0.41 | 0.003 | 0.03±0.133 | 1.63 | 0.19 | 0.154 |
| ***Temporal*** | 0.06±1.094 | 1.38 | 0.06 | 0.169 | 0.02±1.145 | 0.51 | 0.02 | 0.612 |
| Fusiform gyrus | -0.00±0.198 | -0.06 | 0.01 | 0.952 | 0.04±0.176 | 1.79 | 0.20 | 0.126 |
| Inferior temporal gyrus | 0.03±0.16 | 1.86 | 0.21 | 0.126 | 0.03±0.141 | 2.03 | 0.23 | 0.093 |
| Middle temporal gyrus | 0.00±0.15 | 0.07 | 0.01 | 0.952 | -0.02±0.123 | -1.46 | 0.17 | 0.203 |
| Superior temporal gyrus | 0.03±0.163 | 1.79 | 0.21 | 0.126 | -0.03±0.139 | -1.70 | 0.20 | 0.144 |

| **B.** | ***Left SP thickness*** | | | | ***Right SP thickness*** | | | |
|---|---|---|---|---|---|---|---|---|
| | **CHD** | **TD** | **D** | **q** | **CHD** | **TD** | **D** | **q** |
| **Global** | -0.013±0.970 | -6.86 | 0.14 | <0.001 | 0.01±0.947 | 0.32 | 0.01 | 0.750 |
| ***Frontal*** | -0.07±1.005 | -1.45 | 0.06 | 0.147 | 0.04±0.985 | 1.03 | 0.04 | 0.305 |
| Middle frontal gyrus | -0.03±0.104 | -2.71 | 0.31 | 0.018 | 0.01±0.093 | 0.87 | 0.10 | 0.526 |
| Precentral gyrus | -0.00±0.089 | -0.45 | 0.05 | 0.768 | 0.03±0.063 | 4.31 | 0.49 | <0.001 |
| Superior frontal gyrus | -0.02±0.083 | -2.34 | 0.27 | 0.039 | -0.0±0.106 | -0.04 | 0.00 | 0.967 |
| Paracentral lobule | -0.03±0.094 | -2.70 | 0.31 | 0.018 | -0.01±0.089 | -0.64 | 0.07 | 0.638 |
| Inferior frontal gyrus | 0.04±0.073 | 4.91 | 0.56 | <0.001 | -0.0±0.082 | -0.17 | 0.02 | 0.892 |
| ***Parietal*** | -0.08±0.967 | -2.37 | 0.09 | 0.018 | 0.10±0.870 | 3.40 | 0.13 | 0.001 |
| Inferior parietal cortex | -0.04±0.089 | -4.17 | 0.48 | <0.001 | 0.03±0.067 | 4.45 | 0.51 | <0.001 |
| Precuneus cortex | -0.05±0.081 | -5.22 | 0.60 | <0.001 | -0.03±0.078 | -3.16 | 0.36 | 0.006 |
| Superior parietal cortex | -0.01±0.112 | -0.68 | 0.08 | 0.630 | 0.05±0.078 | 5.19 | 0.60 | <0.001 |
| Supramarginal gyrus | 0.00±0.089 | 0.25 | 0.03 | 0.861 | 0.03±0.086 | 3.51 | 0.40 | 0.002 |
| Postcentral gyrus | 0.02±0.091 | 1.54 | 0.18 | 0.198 | 0.03±0.078 | 3.01 | 0.35 | 0.009 |
| ***Occipital*** | -0.26±0.934 | -5.92 | 0.29 | <0.001 | -0.06±0.984 | -1.39 | 0.07 | 0.167 |
| Cuneus | -0.09±0.105 | -7.79 | 0.89 | <0.001 | -0.03±0.103 | -2.50 | 0.29 | 0.028 |
| Lateral occipital cortex | -0.02±0.086 | -2.32 | 0.27 | 0.040 | 0.01±0.092 | 0.84 | 0.10 | 0.526 |
| Lingual gyrus | -0.10±0.123 | -6.88 | 0.79 | <0.001 | -0.02±0.145 | -1.45 | 0.17 | 0.224 |
| ***Temporal*** | -0.15±0.948 | -3.96 | 0.07 | <0.001 | -0.09±0.951 | -2.41 | 0.10 | 0.016 |
| Fusiform gyrus | -0.06±0.104 | -5.16 | 0.59 | <0.001 | -0.04±0.109 | -3.15 | 0.36 | 0.006 |
| Inferior temporal gyrus | 0.00±0.081 | 0.38 | 0.04 | 0.801 | -0.01±0.111 | -1.16 | 0.13 | 0.355 |
| Middle temporal gyrus | -0.04±0.08 | -4.45 | 0.51 | <0.001 | -0.03±0.097 | -2.53 | 0.29 | 0.027 |
| Superior temporal gyrus | -0.01±0.067 | -1.84 | 0.21 | 0.114 | 0.00±0.085 | 0.24 | 0.03 | 0.861 |

**A. SP volume**

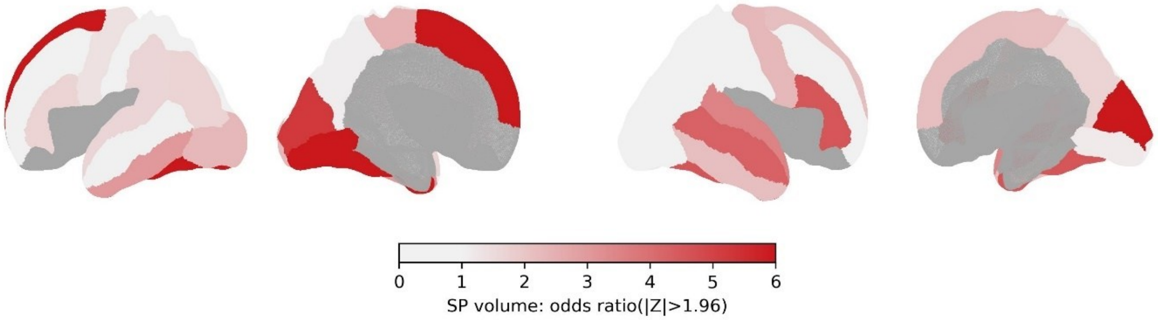


**B. SP thickness**

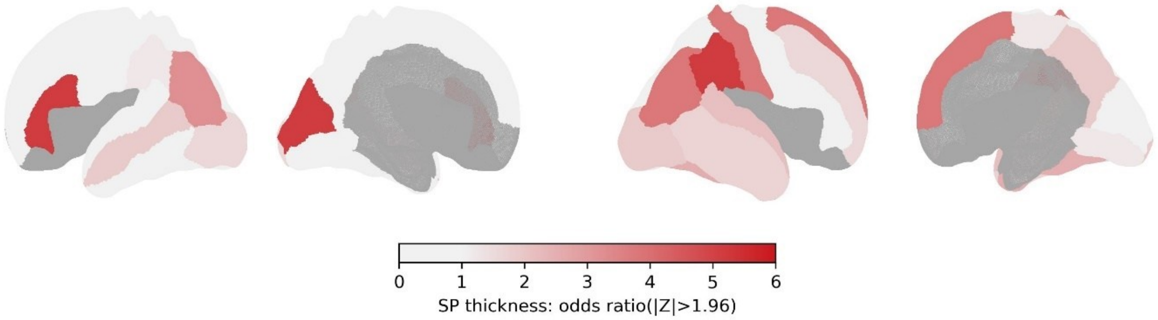


**SI-Fig11.** Comparison of extreme deviations in CHD subjects relative to TD controls for SP volume (**A**) and SP thickness (**B**). Surface maps display regional odds ratios indicating the likelihood of |Z| > 1.96 with values >1 indicating higher probability.

**SI-Table7A.** Type II analysis of variance for the linear models evaluating relationships between SP deviations and substrate delivery scores and CPR (Z-scores) (including Echo-MRI delay as a covariate) independently, and in joint models. Interaction between substrate delivery scores and CPR were tested only for SP thickness deviations as neither of the two measures reached significance in SP volume deviation models. Reported statistics include F-statistics, p-values (p), and partial eta-squared ($\eta^2$). For significant associations, we also report the slope of the relationship ($\beta$) **B.** SP thickness deviations split across ventricle physiology.

**A.**

| | | F | p | $\eta^2$ | β |
|---|---|---|---|---|---|
| Volume ratio (GA, sex) | Substrate delivery score | 2.45 | 0.112 | 0.03 | |
| | Cerebroplacental ratio (Z) | 0.04 | 0.851 | 0.00 | |
| | Echo-MRI delay (weeks) | 0.78 | 0.382 | 0.01 | |
| | Substrate delivery score | 1.71 | 0.195 | 0.03 | |
| | Cerebroplacental ratio (Z) | 0.11 | 0.737 | 0.00 | |
| | Echo-MRI delay (weeks) | 0.66 | 0.421 | 0.01 | |
| Volume ratio (GA, sex, brain size) | Substrate delivery score | 1.67 | 0.200 | 0.02 | |
| | Cerebroplacental ratio (Z) | 3.14 | 0.081 | 0.06 | |
| | Echo-MRI delay (weeks) | 0.26 | 0.613 | 0.00 | |
| | Substrate delivery score | 2.40 | 0.126 | 0.04 | |
| | Cerebroplacental ratio (Z) | 3.82 | 0.055 | 0.06 | -0.55 |
| | Echo-MRI delay (weeks) | 0.18 | 0.673 | 0.00 | |
| Thickness ratio (GA, sex) | Substrate delivery score | 2.19 | 0.143 | 0.03 | |
| | Cerebroplacental ratio (Z) | 3.61 | 0.062 | 0.05 | -0.53 |
| | Echo-MRI delay (weeks) | 0.08 | 0.780 | 0.00 | |
| | Substrate delivery score | 3.57 | 0.063 | 0.05 | -0.23 |
| | Cerebroplacental ratio (Z) | 4.58 | 0.036 | 0.07 | -0.59 |
| | Echo-MRI delay (weeks) | 0.03 | 0.860 | 0.00 | |
| Thickness ratio (GA, sex, brain size) | Substrate delivery score | 1.07 | 0.305 | 0.01 | |
| | Cerebroplacental ratio (Z) | 6.06 | 0.017 | 0.08 | -0.65 |
| | Echo-MRI delay (weeks) | 0.01 | 0.927 | 0.00 | |
| | Substrate delivery score | 2.73 | 0.103 | 0.04 | |
| | Cerebroplacental ratio (Z) | 7.07 | 0.010 | 0.10 | -0.59 |
| | Echo-MRI delay (weeks) | 0.04 | 0.852 | 0.00 | |

**B.**

| | | Single ventricle (deviation mean ± SD) | Two ventricle (deviation mean ± SD) | β | p (q) |
|---|---|---|---|---|---|
| Middle frontal gyrus | lh | -0.07±0.122 | -0.02±0.092 | -0.06 | 0.026 (0.220) |
| Precentral gyrus | rh | -0.03±0.071 | 0.01±0.095 | -0.04 | 0.020 (0.220) |
| Superior frontal gyrus | rh | 0.01±0.062 | 0.04±0.062 | -0.07 | 0.013 (0.220) |
| Superior frontal gyrus | lh | -0.06±0.109 | -0.01±0.066 | -0.05 | 0.017 (0.220) |
| Precentral gyrus | lh | -0.05±0.165 | 0.02±0.063 | -0.04 | 0.074 (0.500) |

**A.**

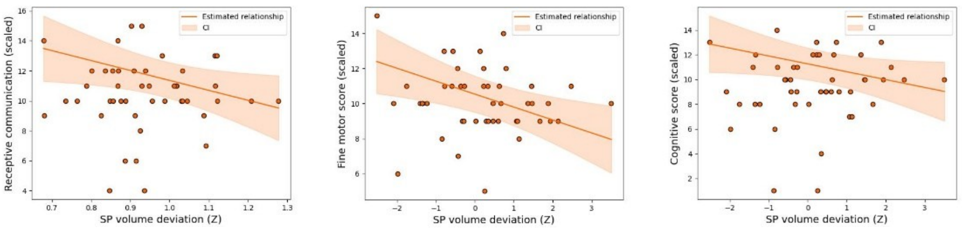


**B.**

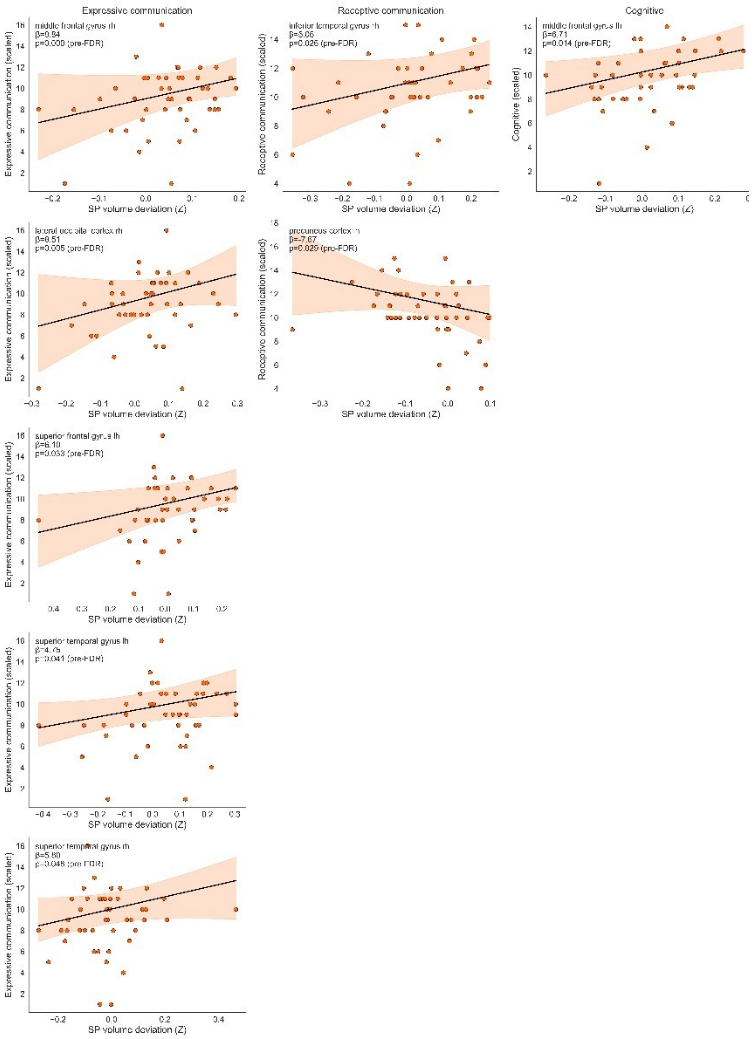
Expressive communication
Receptive communication
Cognitive
middle frontal gyrus rh
β=0.84
p=0.009 (pre-FDR)
inferior temporal gyrus rh
β=5.06
p=0.026 (pre-FDR)
middle frontal gyrus lh
β=6.71
p=0.014 (pre-FDR)
lateral occipital cortex rh
β=8.51
p=0.005 (pre-FDR)
precuneus cortex rh
β=-7.67
p=0.029 (pre-FDR)
superior frontal gyrus lh
β=6.10
p=0.033 (pre-FDR)
superior temporal gyrus lh
β=4.75
p=0.041 (pre-FDR)
superior temporal gyrus rh
β=5.60
p=0.048 (pre-FDR)
Expressive communication (scaled)
Receptive communication (scaled)
Cognitive (scaled)
SP volume deviation (Z)

**C.**

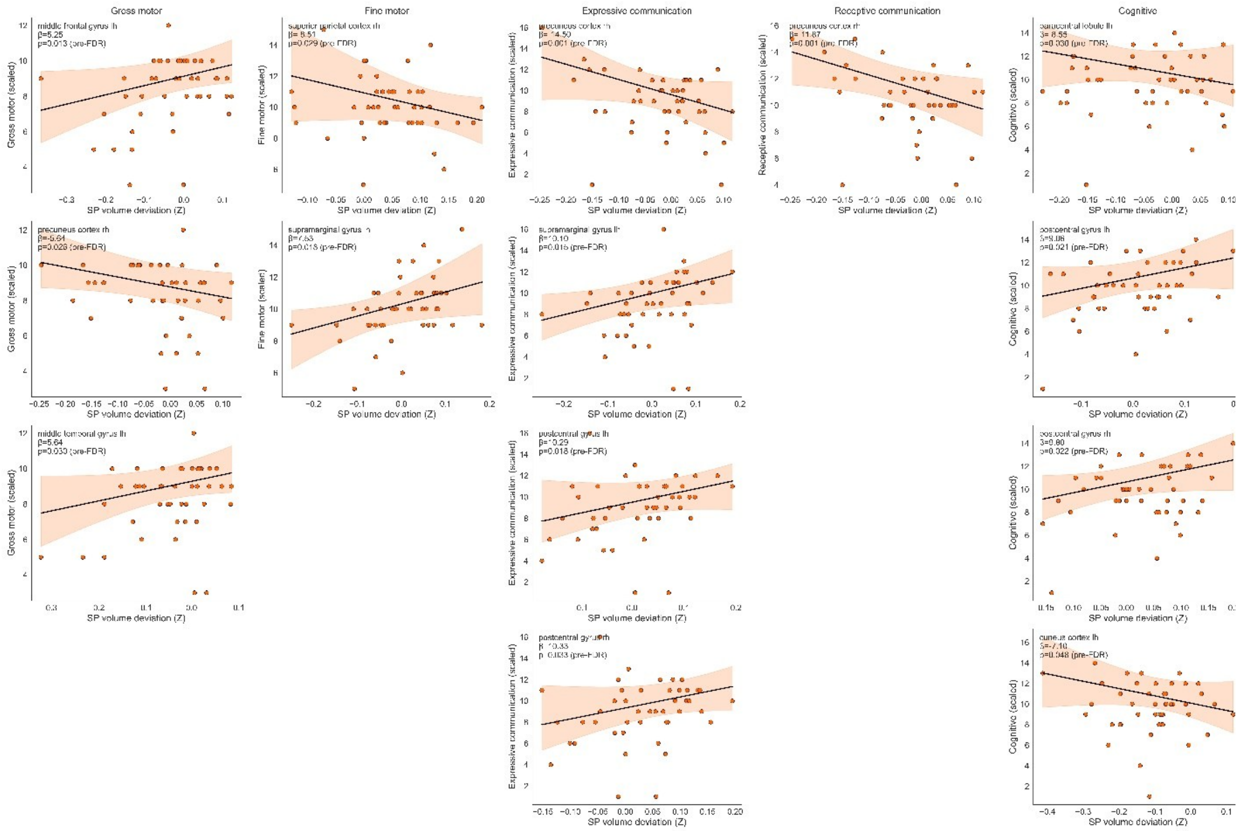

**SI-Fig12.** Relationships of regional SP deviations with neurodevelopmental outcomes. **A.** Scatter plot showing the association between whole-brain SP volume deviations and fine motor outcomes: SP volume ratio (GA, sex TD-referenced) on the left, SP volume ration (GA, sex, whole-brain volume TD-rederenced) in the middle and the right. **B.** Scatter plots of regional SP volume deviations versus developmental scores. No regions remained significant after FDR correction; only regions with pre-FDR significance are shown for illustrative purposes. Plots are ordered as score per column. **C.** Same as B but for SP thickness deviations. In all plots, each point represents a participant; regression lines indicate the direction and magnitude of the relationship estimated by robust regression.

**SI-Table8.** Results of SP deviation vs neurodevelopmental outcome comparisons reporting slopes of relationships estimated with robust linear models, pre-FDR p-value, and post-FDR q value.

***SP volume deviation***

| *Outcome* | *Region* | β | p | q |
|---|---|---|---|---|
| **Receptive communication** | inferior temporal gyrus_rh | 5.06 | 0.026 | 0.493 |
| | precuneus cortex_lh | -7.67 | 0.029 | 0.493 |
| **Expressive communication** | middle frontal gyrus_rh | 9.84 | 0.009 | 0.153 |
| | lateral occipital cortex_rh | 8.51 | 0.005 | 0.153 |
| | superior frontal gyrus_lh | 6.10 | 0.033 | 0.241 |
| | superior temporal gyrus_lh | 4.75 | 0.041 | 0.241 |
| | superior temporal gyrus_rh | 5.80 | 0.048 | 0.241 |

| | | | | |
|---|---|---|---|---|
| **Cognitive** | middle frontal gyrus_lh | 6.71 | 0.014 | 0.473 |

***SP thickness deviation***

| *Outcome* | *Region* | **β** | **p** | **q** |
|---|---|---|---|---|
| **Gross motor** | middle frontal gyrus_lh | 5.25 | 0.013 | 0.342 |
| | precuneus cortex_rh | -5.64 | 0.026 | 0.342 |
| | middle temporal gyrus_lh | 5.64 | 0.030 | 0.342 |
| **Fine motor** | superior parietal cortex_rh | -8.51 | 0.029 | 0.488 |
| | supramarginal gyrus_lh | 7.53 | 0.018 | 0.488 |
| **Receptive communication** | precuneus cortex_rh | -11.87 | 0.001 | 0.017 |
| **Expressive communication** | precuneus cortex_rh | -14.50 | 0.001 | 0.039 |
| | supramarginal gyrus_lh | 10.10 | 0.015 | 0.203 |
| | postcentral gyrus_lh | 10.29 | 0.018 | 0.203 |
| | postcentral gyrus_rh | 10.33 | 0.033 | 0.280 |
| **Cognitive** | paracentral lobule_lh | -8.55 | 0.030 | 0.344 |
| | postcentral gyrus_lh | 9.06 | 0.021 | 0.344 |
| | postcentral gyrus_rh | 9.80 | 0.022 | 0.344 |
| | cuneus cortex_lh | -7.10 | 0.048 | 0.366 |
